# Pharmacologic DPP-4 inhibition promotes CD8⁺ T cell metabolic fitness to enhance anti-tumor activity

**DOI:** 10.64898/2026.03.31.715681

**Authors:** Oriana Y. Teran Pumar, Elton L VanNoy, Abigail Haffey, Durga Prasad Gannamedi, Christine Isabelle Rafie, Dylan Scott Lykke Harwood, Julia R. Benedetti, Christine Ann Pittman Ballard, Erika Ciervo, Pedro Henrique Assenza Tavares Coroa, Payal Grover, Brandon Emanuel León, Jonathan Mitchell, Asmita Pathak, Bruno Colon, Lyenne El Ghorayeb, Laura O’Sullivan, Venu Venkatarame Gowda Saralamma, Clara Lopez Ruiz, Natasha Khatwani, Surinder Kumar, Priyamvada Rai, Jonathan Schatz, Ashish Shah, Zev A. Binder, Michele Ceccarelli, Quinn T. Ostrom, Bjarne Winther Kristensen, Erietta Stelekati, Dionysios C. Watson, David B. Lombard, Dalia Haydar, Defne Bayik

## Abstract

Metabolic dysfunction is a hallmark of CD8^+^ T cell exhaustion in the tumor microenvironment. Thus, there is growing interest in developing strategies that enhance anti-tumor functions of CD8^+^ T cells via metabolic reprogramming. Here, we identify dipeptidyl peptidase 4 (DPP-4) as a previously unknown regulator of CD8^+^ T cell function and metabolism. We discovered that DPP-4 is upregulated in exhausted CD8^+^ T cells. Pharmacological inhibition of DPP-4 with the FDA-approved anti-diabetic drug sitagliptin transcriptionally and metabolically reprogrammed CD8^+^ T cells, increasing spare mitochondrial respiratory capacity, proliferation, cytotoxic mediator production, and antigen-specific cancer cell killing capability. The functional effects of sitagliptin were dependent on upregulation of glutamate decarboxylase 1 (GAD1), an enzyme that feeds glutamate into the tricarboxylic acid (TCA) cycle, highlighting a new role for GAD1 in CD8^+^ T cell respiration and proliferation. We found that systemic inhibition of DPP-4 in preclinical mouse glioblastoma (GBM) models prolongs survival in a CD8^+^ T cell-dependent manner, and retrospective clinical cohort analysis revealed better outcomes in GBM patients using DPP-4 inhibitors. Importantly, preconditioning of Chimeric Antigen Receptor (CAR) T-cells with DPP-4 inhibition enhanced their cytotoxicity, persistence, and therapeutic efficacy in pediatric GBM. Together, our findings provide mechanistic and biological rationale for repurposing readily accessible DPP-4 inhibitors to enhance anti-tumor CD8^+^ T cell responses.

## Introduction

Cancer therapy has been revolutionized by the discovery and therapeutic targeting of immune checkpoints that suppress the anti-tumor function of T cells^1,2^. Immune checkpoint signaling is a hallmark of exhausted T cells with lower effector potential driven by an immunosuppressive tumor microenvironment (TME)^3–5^. Increasing evidence demonstrates that T cell exhaustion is tightly linked to metabolic dysfunction, which impairs overall anti-tumor activity of host T cells and limits the therapeutic efficacy of Chimeric Antigen Receptor (CAR) T-cells^6–15^. Thus, modulating T cell metabolism is a promising and underexplored strategy to potentiate anti-tumor immune response, particularly in cancers where immunotherapies have had limited efficacy, such as glioblastoma (GBM) - the most common primary malignant brain tumor^16–18^.

CD8^+^ T cells metabolism is dynamically regulated throughout the maturation and activation states. Naïve CD8^+^ T cells, which are metabolically quiescent, primarily rely on basal levels of oxidative phosphorylation (OXPHOS)^19–21^. T cell activation, upon cognate antigen recognition to elicit an immune response, depends on a metabolic rewiring that elevates both mitochondrial respiration and glycolysis^22,23^. After differentiation into the effector state, T cells also enhance fatty acid synthesis and amino acid/nucleotide metabolism to compensate for the bioenergetic demands of continued proliferation and cytotoxic activity^22,24–29^. However, as CD8^+^ T cells become exhausted, they experience another metabolic shift characterized by a decline in mitochondrial activity^30^ and overreliance on glycolysis^31^. Decreased mitochondrial membrane potential and increased oxidative stress are key features of this metabolic phenotype and are closely associated with T cell dysfunction^6,11,32^. Given the tight link between cellular metabolism and immune function, metabolic reprogramming of CD8⁺ T cells has been explored in preclinical models of cancer^8,9,24,33–40^. However, identification of metabolic targets with a viable therapeutic window remains a major obstacle for clinical translation of these approaches in the context of cancer.

We previously observed that dipeptidyl peptidase IV (DPP-4, CD26) is expressed by a subset of immunosuppressive myeloid cells in GBM^41^. DPP-4 is a transmembrane and soluble exopeptidase widely studied for its role in regulating whole-body glucose metabolism via inactivation of glucagon-like peptide-1 (GLP-1)^42,43^. Due to this central role, DPP-4 inhibitors, named gliptins, have been developed to pharmacologically block DPP-4 enzymatic activity and approved by the FDA/EMA as anti-diabetic agents^42,44^. DPP-4 can also play a role in systemic immunoregulation by cleaving cytokines/chemokines to regulate their levels and bioactivity^45–47^. As such, DPP-4 inhibition was shown to indirectly drive CD8^+^ T cell tumor infiltration by enhancing CXCL10 levels in preclinical models of peripheral cancers^48–50^. However, cellular roles of DPP-4 remain poorly defined. In this study, we identify DPP-4 is a direct modulator of CD8^+^ T cell activity, and metabolism and demonstrate that it serves as an immunotherapy target in GBM. DPP-4 was a marker of CD8^+^ T cell exhaustion, and its pharmacological inhibition enhanced cytotoxic function of CD8^+^ T cells. We observed that DPP-4 inhibitor use is associated with better overall survival of GBM patients and DPP-4 inhibition improved GBM outcomes in preclinical models in a manner dependent on CD8^+^ T cells. Our results identified glutamate decarboxylase (GAD1) as a critical downstream effector driving enhanced metabolic fitness of CD8^+^ T cells upon DPP-4 inhibition. DPP-4 inhibition also enhanced persistence and therapeutic efficacy of CAR T-cells directed toward pediatric GBM. Thus, our study establishes DPP-4 as a therapeutically relevant CD8^+^ T cell checkpoint molecule that can be targeted by repurposing accessible and safe drugs to improve cancer immunotherapy responses.

## Results

### DPP-4 is a CD8^+^ T cell-associated immunotherapy target in GBM

Our earlier studies suggested that systemic inhibition of DPP-4 extends survival in preclinical models of GBM^41^. Thus, we aimed to determine whether DPP-4 inhibition could comprise a viable anti-tumor strategy in the clinic. We used data from The National Cancer Institute’s Surveillance, Epidemiology, and End Results (SEER) program linked to Medicare claims to retrospectively assess the survival probability in primary GBM patients receiving standard therapy with surgical resection, radiation, and temozolomide alone (Standard of Care, SOC) or in combination with either metformin (an irrelevant anti-diabetic drug; Metformin+SOC) or DPP-4 inhibitors (DPP4i+SOC). Median survival in the SOC only group was 12.4 months (95% CI: 12.0-13.0), compared to 13 months (95% CI: 11.8-14.7) in the Metformin+SOC group, and 15.9 months (95% CI: 10.0-31.0, p-value=0.41) in the DPP4i+SOC group. Indeed, 19 individuals who received DPP-4i experienced a 47% decrease in risk of death as compared to those that received SOC alone (HR: 0.63, 95% CI: 0.38-1.04, p-value=0.070; **Fig. 1A, Fig. S1**). To determine the mechanism of DPP-4 inhibitor response, we assessed DPP-4 surface expression in a human glioma single cell dataset (GSE163108^51^) and discovered that *DPP4* was primarily expressed by lymphocytes and a subset of cancer cells (**Fig. S2A-C**). Among the lymphocyte population, several CD8^+^ T cell populations were characterized by high *DPP4* expression (**Fig. S2D**). These results were supported by a second dataset, (GSE117891^52^), where we observed that tumor-infiltrating CD8^+^ T cells had higher levels of *DPP4* expression compared to other immune populations (**Fig. 1B**). Matching the patient data, CD8^+^ T cells infiltrating mouse GBM expressed the highest levels of DPP-4 among the defined immune lineages, a pattern conserved in spleens (**Fig. 1C-D**, **S3A-E**). Thus, to assess the role of CD8^+^ T cells in DPP-4 inhibitor-mediated survival extension, we administered sitagliptin via drinking water to SB28-implanted mice and concurrently depleted CD8^+^ T cells (**Fig. S4A**). While sitagliptin administration conferred a survival advantage, this effect was not observed when CD8^+^ T cells were depleted, with median survival remaining indistinguishable between sitagliptin- and vehicle-treated mice (**Fig. S4B**). Correspondingly, sitagliptin had no direct effect on the viability of cancer cells (**Fig. S4C**). Collectively, these findings suggest CD8^+^ T cell-associated DPP-4 serves an immunotherapy target in GBM.

**Figure 1.**
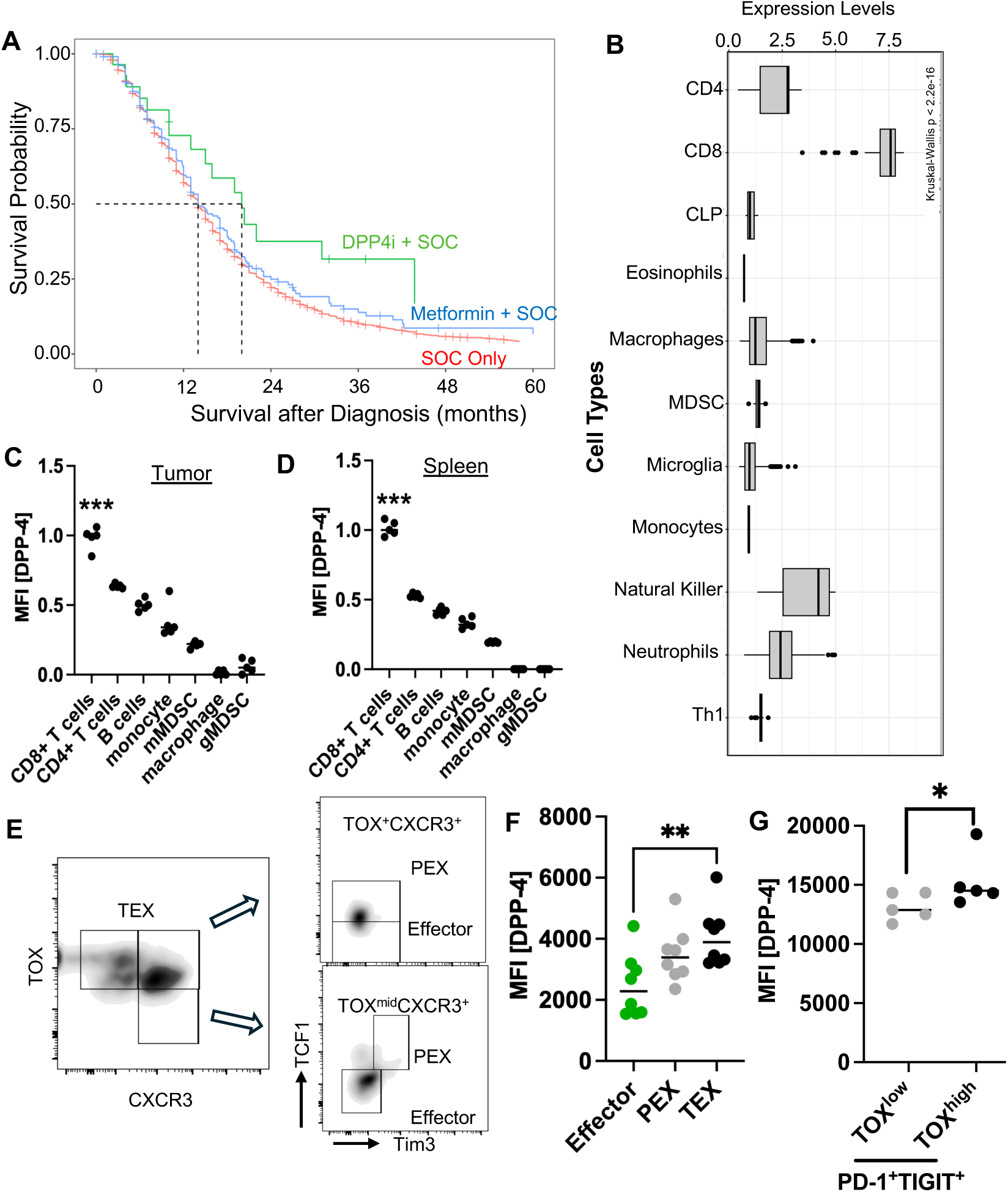
DPP4 inhibitors improve GBM outcomes by targeting CD8+ T-Cells. **(A)** Kaplan-Meier curve of GBM patients on standard of care (SOC; red, median=12.4 months) or combinations of SCO with metformin (blue, media=13 months) or DPP4 inhibitors (DPP4i, median=15.9 months) based on the SEER dataset. **(B)** *DPP4* expression from immune lineages as determined by the scRNA-seq of 13 brain tumour patients (3 Grade II; 1 Grade III-IV; 9 GBM). Mean fluorescence intensity (MFI) of DPP-4 from **(C)** tumor-infiltrating and **(D)** splenic immune populations. n=5; *** p<0.001 by two-way ANOVA. **(E)** Representative gating strategy used to define effector, progenitor exhausted (PEX), or terminally exhausted (TEX) populations. **(F)** DPP-4 MFI from effector, PEX, and TEX populations. n=8; ** p<0.01 by two-way ANOVA. **(G)** Exhausted CD8^+^ T cells from GBM and tumor-adjacent tissue were defined as: CD45^+^CD11b^-^CD3^+^PD1^+^TIGIT^+^. DPP-4 MFI from TOX^+^ and TOX^-^population. n=4; * p<0.05 by paired t-test.

### DPP-4 expression on CD8^+^ T cells is informed by the activation state

Since earlier studies demonstrated that DPP-4 was upregulated in memory CD4^+^ T cells^53,54^, we interrogated whether CD8^+^ T cell activation status informs DPP-4 levels using PBMCs from healthy human donors. Similar to CD4^+^ T cells, effector CD8^+^ T cells had lower DPP-4 expression compared to memory lineages (**Fig. S5A-B**). These observations were further confirmed by using an acute LCMV infection model, where memory CD8^+^ T cells had significantly higher DPP-4 levels compared to the effector subset (**Fig. S5C-D**). Since memory and exhausted T cells have some overlapping phenotypes^55,56^, we also interrogated whether DPP-4 expression profile was informed by exhaustion. In the chronic LCMV infection model, we observed that exhausted T cells had increased DPP-4 levels compared to effector cells (**Fig. S5C-D**). To evaluate dynamic changes in DPP-4 expression, we performed an *in vitro* CD8^+^ T cell exhaustion assay. CD8^+^ T cells from OT-I mice were either activated for two days or exhausted by repeated OVA stimulation for 6 days (**Fig. S5E**). We observed a gradual increase in DPP-4 with terminally exhausted T cells (TEX) having significantly higher DPP-4 levels compared to the effector subset (**Fig. 1E-F**). Finally, to further confirm the correlation between DPP-4 expression and CD8^+^ T cell exhaustion phenotype, we analyzed high-grade glioma samples and tumor-adjacent tissue (**Fig. S5F**). We discovered that PD1^+^TIGIT^+^TOX^+^ CD8^+^ T cells, marking the TEX phenotype, had significantly higher DPP-4 expression compared to PD1^+^TIGIT^+^TOX^-^ CD8^+^ T cells (**Fig. 1G**). Collectively, these findings suggest that DPP-4 is upregulated in exhausted CD8^+^ T cells, including in GBM.

### DPP-4 inhibition promotes CD8^+^ T cell effector function

Given that DPP-4 expression increased in exhausted CD8^+^ T cells, we sought to determine whether DPP-4 functioned as an immune checkpoint and if its inhibition could be used to promote effector activity. We activated splenic CD8^+^ T cells with anti-CD3/CD28 plus IL-2 for 4 days in the presence of the DPP-4 inhibitors sitagliptin or saxagliptin (**Fig. 2A**). Pharmacological inhibition of DPP-4 significantly increased T cell proliferation (**Fig. 2B**) and maintained proliferative capacity as observed by enhancement of the Ki-67^+^ population (**Fig. S6A**). Similarly, sitagliptin enhanced the proliferation of CD8+ T cells obtained from the PBMCs of healthy donors (**Fig. S6B**). To determine whether this activation phenotype translated into effector function, we assessed the production of cytotoxic mediators Granzyme B (GzmB) and interferon gamma (IFNγ) from OT-I cells under exhaustion conditions. There was a significant increase in expression of both cytotoxic mediators in the presence of sitagliptin (**Fig. 2C-D**). We also performed an antigen-specific cancer cell killing assay. CD8^+^ T cell from OT-I mice were activated for two days in the presence of sitagliptin or vehicle control, and then co-cultured with OVA-expressing or parental SB28 at a 1:1 ratio for 6 hours (**Fig. S6C**). We observed that OVA-expressing cancer cells experienced higher cell death rates when co-cultured with OT-I T cells that had been activated in the presence of sitagliptin (**Fig. 2F-G**). In contrast, nonspecific killing of parental SB28 cells was not impacted by sitagliptin treatment of CD8^+^ T cells. This overall enhanced effector function led us to evaluate whether DPP-4 inhibition could also overcome exhaustion. In the setting of sustained stimulation of wild-type T cells *in vitro* or chronic LCMV infection *in vivo*, sitagliptin treatment resulted in a reduction in the frequency of T cells co-expressing the inhibitory receptors PD-1 and 2B4 (**Fig. S6D-F**). There was also an increase in the frequency of effector CD8^+^ T cell population with sitagliptin treatment under chronic OVA stimulation of OT-I T cells (**Fig. S6G**), suggesting that DPP-4 inhibition enhances CD8^+^ T cell function and dampens T cell exhaustion.

**Figure 2.**
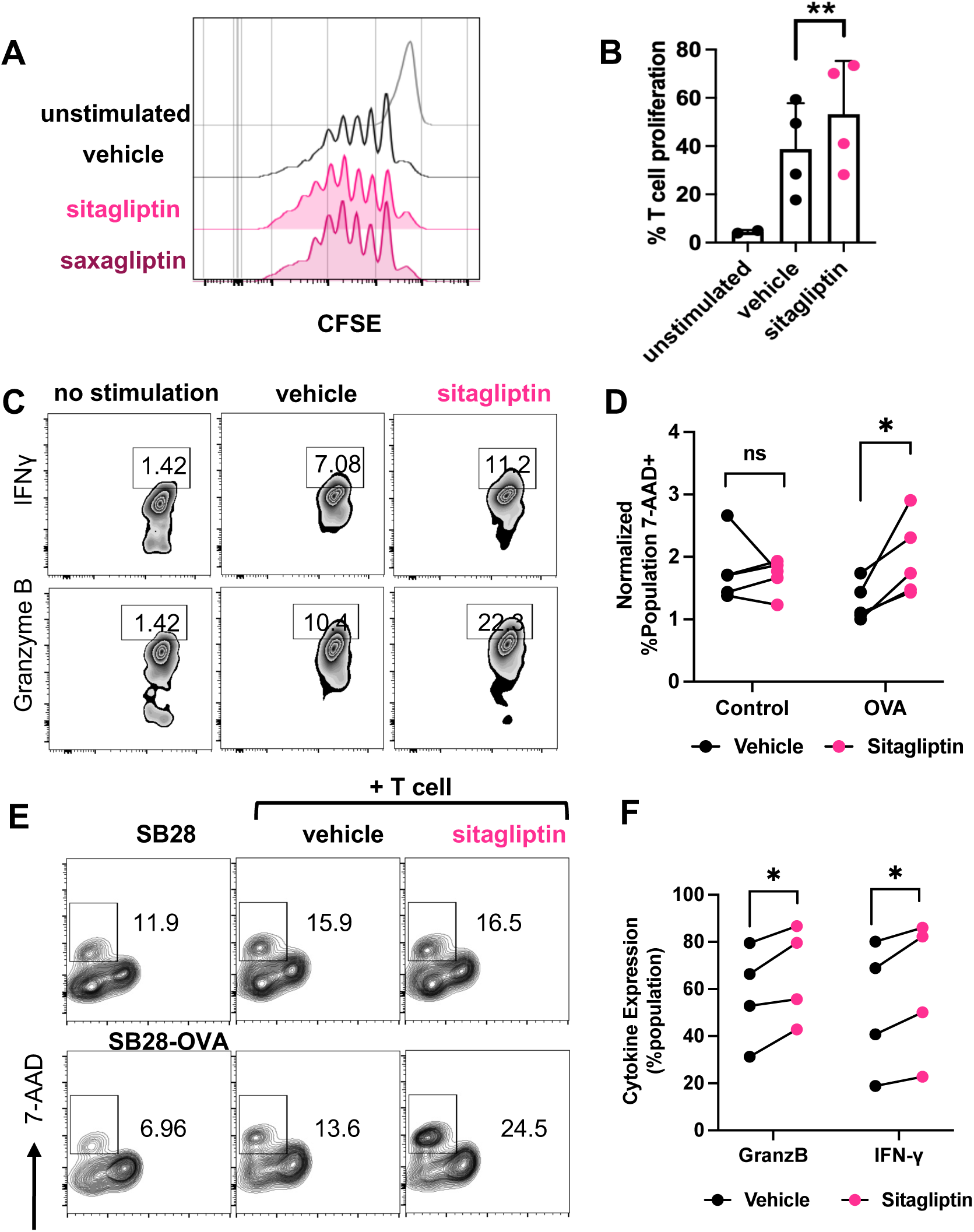
DPP-4 inhibition supports T cell immunologic function. **(A)** Representative histogram depicting CFSE dilution of mouse CD8^+^ T cells activated with anti-CD3/CD28 plus 100 IU/mL IL-2 in the presence of 100 μM sitagliptin, saxagliptin, or vehicle control for 4 days. **(B)** The percentage of proliferating mouse CD8^+^ T cells treated with sitagliptin vs. control. n=4; ** p<0.01 by paired t-test. **(C)** Representative zebra plots depicting Granzyme B and IFNγ production from OT-I T cells on Day 5. **(D)** The percentage of OT-I T cells expressing high levels of Granzyme B or IFNγ. n=4; * p<0.05 by paired t-test. **(E)** Representative zebra plot showing 7-AAD levels in SB28 and ovalbumin-expressing SB28 (SB28-OVA) cells co-cultured with OT-I CD8^+^ T cells at a 1:1 ratio. **(F)** The frequency of lysed wild-type SB28 or SB28-OVA cells cultured with sitagliptin- or vehicle-treated OT-I T cells. n=5; * p<0.05, ns=not significant by paired t-test.

### Sitagliptin treatment transcriptionally and metabolically reprograms CD8^+^ T cells

To interrogate the mechanisms by which DPP-4 informed T cell function, we initially performed transcriptional analysis. DPP-4 inhibition significantly upregulated pathways associated with lymphocyte activation, whereas there was no consistent pattern for downregulated pathways (**Fig. 3A, S7A-B**). In addition, mitochondria-related pathways were enriched in sitagliptin-treated T cells compared to vehicle controls (**Fig. 3A**). This prompted us to evaluate potential differences in cellular bioenergetics. CD8^+^ T cells activated in the presence of sitagliptin had higher rates of basal and maximal respiratory capacity (**Fig. 3B-C**), consistent with increased OXPHOS observed in activated T cells^25,28^. While mitogenesis was comparable in both groups (**Fig. S7C-D**), we detected higher mitochondrial polarization, enhanced proton efflux rate (PER), and increased total ATP in the sitagliptin-treated T cells (**Fig. 3D-E, S7E**). Since uncontrolled OXPHOS could result in overproduction of reactive oxygen species (ROS)^12^, which is closely linked to the dysfunction of exhausted T cells^11^, we measured mitochondrial superoxide levels with MitoSox staining. There was a significant decrease in MitoSox^+^ CD8^+^ T cell frequency suggesting that DPP-4 inhibition does not promote redox signaling in T cells (**Fig. S7F**). We also analyzed changes in glycolysis, as this metabolic pathway is upregulated in effector T cells^31^. Both glycolysis rate and glucose uptake of CD8^+^ T cells were enhanced upon DPP-4 inhibition (**Fig. 3F-G**), pointing to overall activation of CD8^+^ T cell metabolic activity. Previous studies indicated that metabolically exhausted T cells contain depolarized mitochondria^6^. We, therefore, stained chronically stimulated OT-I cells with the membrane potential-sensitive dye TMRM. There was a significant increase in TMRM intensity in sitagliptin-treated cells (**Fig. S7G**), indicating that DPP-4 inhibition mitigated the characteristic mitochondrial dysfunction in exhausted T cells^6,32^. Given these changes in bioenergetics, we next interrogated the differentially regulated metabolic pathways by performing targeted metabolomics. Pathways linked to cellular proliferation, such as nucleotide and amino acid metabolisms, were significantly upregulated upon DPP-4 inhibition (**Fig. 4A**). In addition, the top 10 enriched metabolic pathways included glycolysis and the tricarboxylic acid (TCA) cycle (**Fig. 4A**), in line with observed enhanced glycolysis and OXPHOS rates. Consistent with this metabolic profile, we also observed that TCA cycle genes were differentially regulated between high vs low *DPP4* expressing CD8^+^ T cells in patients with GBM (**Fig. S8A**). Taken together, these observations suggest that DPP-4 inhibition leads to metabolic reprograming that supports proliferation and effector functions of CD8^+^ T cells.

**Figure 3.**
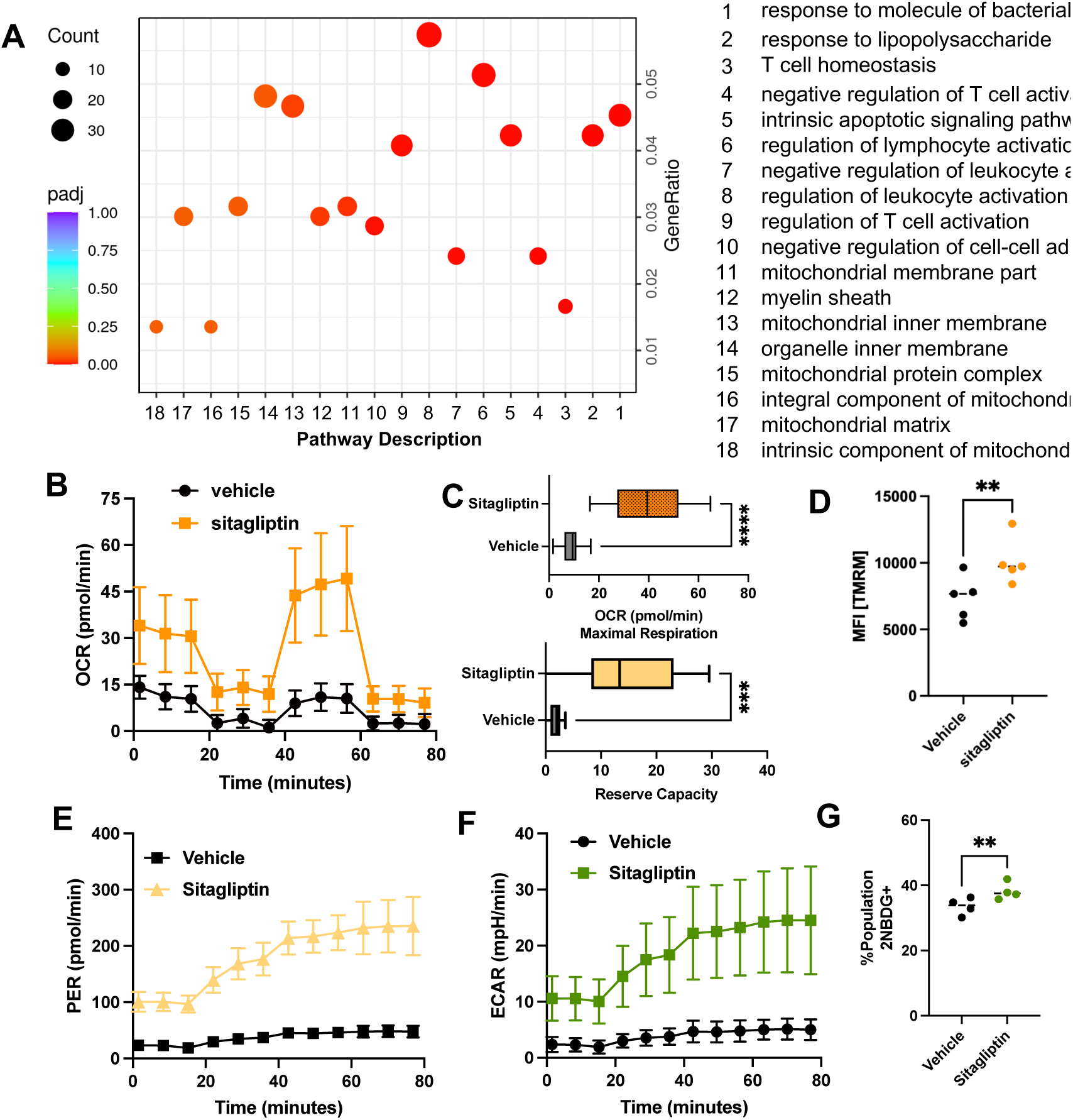
DPP-4 inhibition transcriptionally and metabolically reprograms CD8^+^ T cells. **(A)** Mouse CD8^+^ T cells were activated with anti-CD3/CD28 plus 100 IU/mL IL-2 in the presence of 100 μM sitagliptin, or vehicle control for 4 days. Top differentially up-regulated pathways in sitagliptin-treated CD8^+^ T cells based on genes expressed >2-fold with p<0.05. n=6. Mouse CD8^+^ T cells were activated with anti-CD3/CD28 plus 100 IU/mL IL-2 in the presence of 100 μM sitagliptin, or vehicle control for 4 days and changes in cellular bioenergetics were determined with a Seahorse assay. **(B)** Oxygen consumption rate (OCR) and **(C)** Maximal respiration (top) and reserve capacity (bottom) of sitagliptin- vs. vehicle-treated CD8^+^ T cells. n=6; *** p<0.001, **** p<0.0001 by t-test. **(D)** Mouse CD8^+^ T cells were activated for 2 days and stained with the mitochondrial membrane potential sensitive dye TMRM at 100 nM for 30 minutes. TMRM MFI of vehicle- vs sitagliptin-treated CD8^+^ T cells. n=5; **p<0.01 by paired t-test. **(E)** Proton efflux rate (PER) and **(F)** extracellular acidification rate (ECAR) of CD8^+^ T cells from B, C. n=6. **(G)** CD8^+^ T cells activated for two days were incubated with the glucose mimetic, 2-NBDG (50 μM) for 30 min. The percentage of T cells with 2-NBDG uptake was determined by flow cytometry. n=4; ** p<0.01 by paired t-test.

**Figure 4.**
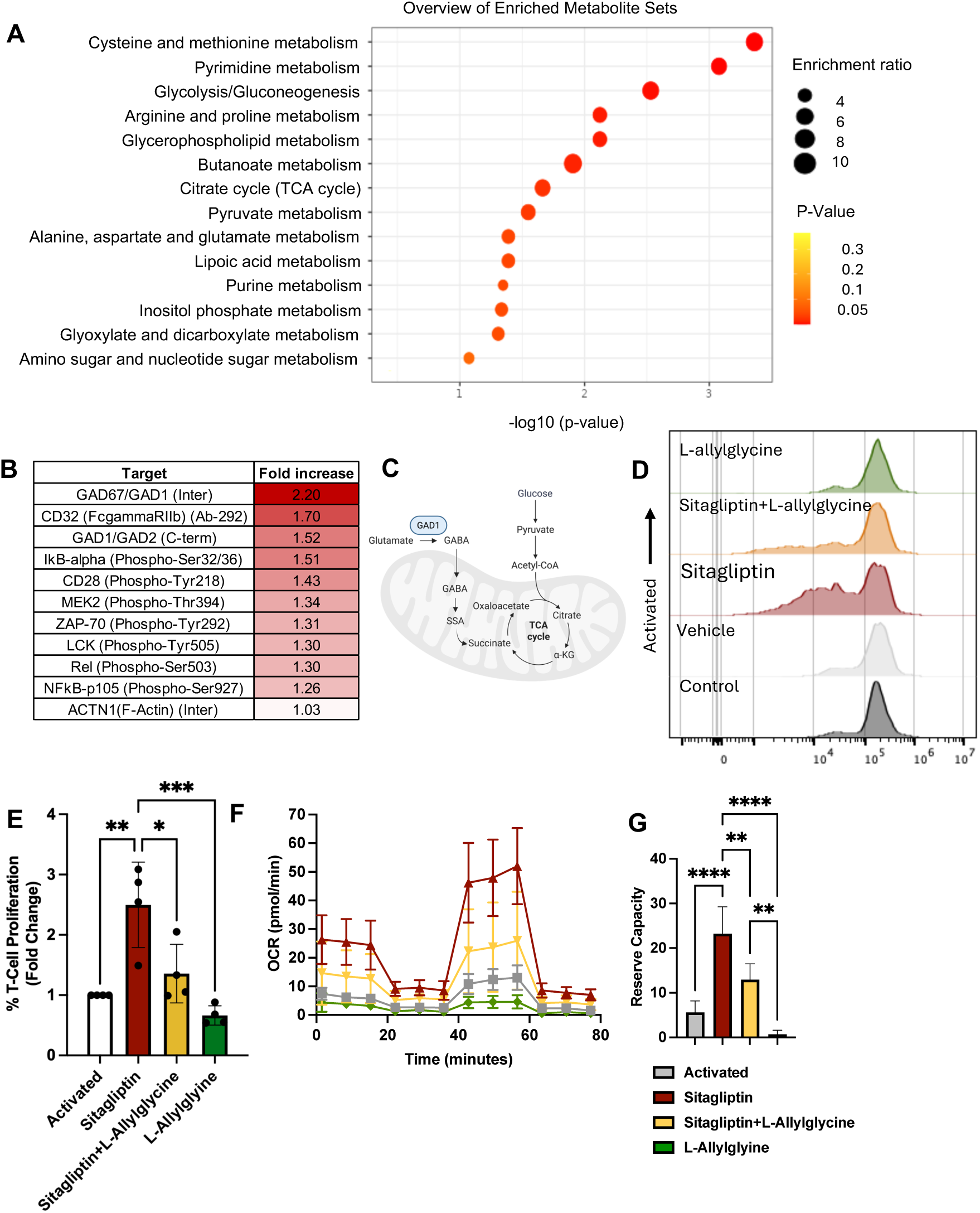
GAD1 is a downstream effector of sitagliptin-mediated T cell reprogramming. **(A)** Targeted metabolomics was performed with mouse CD8^+^ T cells activated for two days with 100 μM sitagliptin or vehicle control. Top enriched metabolic pathways were determined by >1.3-fold increase using Metaboanalyst, n=4. **(B)** T cell receptor (TCR) signaling-focused phosphoproteomics was performed with CD8^+^ T cells activated overnight. Top protein hits that were upregulated or phosphorylated >2-fold in sitagliptin- vs vehicle-treated CD8^+^ T cells. **(C)** Graphical abstract of how GAD1 feeds glutamate to the TCA cycle. **(D)** Representative histograms depicting CFSE dilution of CD8^+^ T cells expanded in the presence of sitagliptin, the GAD1 inhibitor, L-Allylglycine (100 μM), or combination treatment for 6 days. **(E)** Fold change in the frequency of proliferating CD8^+^ T cells normalized to background activation. n=4; * p<0.05, ** p<0.01, *** p<0.001 by one–way ANOVA. **(F)** OCR, and**(G)** reserve capacity of CD8^+^ T cells after 4 days of treatment with sitagliptin, L-Allylglycine (100 μM), or combination treatment, n=6; ** p<0.01, **** p<0.0001 by ordinary one–way ANOVA.

### GAD1 drives sitagliptin-mediated T cell reprogramming

To define the signaling nodes that mediate CD8^+^ T cell reprogramming upon DPP-4 inhibition, we performed a targeted phospho-array screen (**Fig. S8B**). The analysis of the top 10 upregulated and phosphorylated proteins identified changes in TCR downstream signaling, including CD28, ZAP-70, LCK, NF-κB, and CD32^57–60^ (**Fig. 4B**). Surprisingly, the top hit, with a >2-fold upregulation was GAD1, an enzyme that catalyzes the conversion of glutamate to γ-aminobutyric acid (GABA) to fuel into the TCA cycle^61,62^(**Fig. 4B-C**). Correspondingly, these protein phosphorylation changes translated to enrichment of T cell activation/differentiation/signaling along with GABA metabolic processing pathways (**Fig. S8C**). Furthermore, succinic acid and succinic semialdehyde (SSA), the two metabolites produced by the GABA shunt pathway, were upregulated with sitagliptin treatment along with the GAD1 substrate L-glutamic acid (**Fig. 4C, Fig. S8D**). In contrast, the levels of alpha-ketoglutaric acid, the TCA precursor of succinic acid was unchanged, underscoring the functional role of the GABA shunt pathway in sitagliptin-treated CD8^+^ T cells (**Fig. 4C, Fig. S8D**). GAD1 has most extensively been studied in neuronal regulation, where L-allylglycine is used as a functional inhibitor^63^. To test whether GAD1 is required for sitagliptin-mediated functional and metabolic programming of T cells, we first performed a T cell proliferation assay. L-allylglycine significantly dampened proliferation of sitagliptin-treated mouse and human CD8^+^ T cells (**Fig. 4D-E, S7E**). Furthermore, GAD1 inhibition also reversed the increase in maximum cellular respiration and reserve capacity induced by sitagliptin treatment (**Fig. 4F-G, S8F-G**). These observations identify GAD1 as a regulator of CD8^+^ T cell metabolism and function, as well as a downstream effector of DPP-4 inhibition.

### Sitagliptin treatment improves CAR T-cell responses

Treatment of hematological malignancies have been revolutionized by CAR-T cell therapy. However, challenges remain in the manufacturing process as cells become exhausted after prolonged expansion^64,65^. We therefore first tested whether DPP-4 inhibition during manufacturing can potentiate CD19-targeting CAR-T cell responses against OCI-LY1, a human diffuse large B-cell lymphoma (DLBCL) cell line. Indeed, we observed an enhancement in cytolysis, when sitagliptin was introduced during the activation or expansion processes (**Fig. S9A**). These results let us to further explore the ability of DPP-4 inhibition to promote CAR-T cell function in models that focus on the central nervous system. We first treated IL-13Rα2-targeting human CAR T-cells, which are currently in clinical trials for recurrent GBM^66,67^, with sitagliptin. DPP-4 inhibition significantly enhanced cytotoxic activity of IL-13Rα2 CAR T-cells (**Fig. S9B**). We further expanded on these findings by evaluating the effect of DPP-4 inhibition on B7-H3-targeting mouse CAR T-cells, a target that is being evaluated in clinical trials for both recurrent GBM, and pediatric brain tumors,^68^ and designated as a breakthrough therapy by the FDA. Upon repetitive co-culture with cancer cells, sitagliptin-treated CAR T-cells continued to proliferate and eliminate the target cells, while vehicle control stopped exhibiting effector capacity after the second stimulation (**Fig. 5A-D**). Consistently, sitagliptin-treated CAR T-cells exhibited central memory phenotype (**Fig. S9C**), which aids in eliciting a long-lasting and potent anti-tumor response^69,70^. To further determine whether these changes translate as improved treatment response and DPP-4 inhibition can in fact potentiate the therapeutic T cell responses, we intracranially injected the syngeneic pediatric glioma line, KAPP.ff, which naturally expresses B7-H3, into immunocompetent mice. After confirmation of tumor formation, the mice were treated with a single therapeutic dose of vehicle or sitagliptin pretreated CAR-T cells on Day 10. Sitagliptin conditioned CAR-T cells achieved a significantly stronger tumor control (**Fig. 5E, S9D-F**). Collectively, these results indicate that DPP-4 inhibition improves immunotherapy responses by supporting the anti-tumor activity of CD8^+^ T cells (**Fig. 5F**).

**Figure 5.**
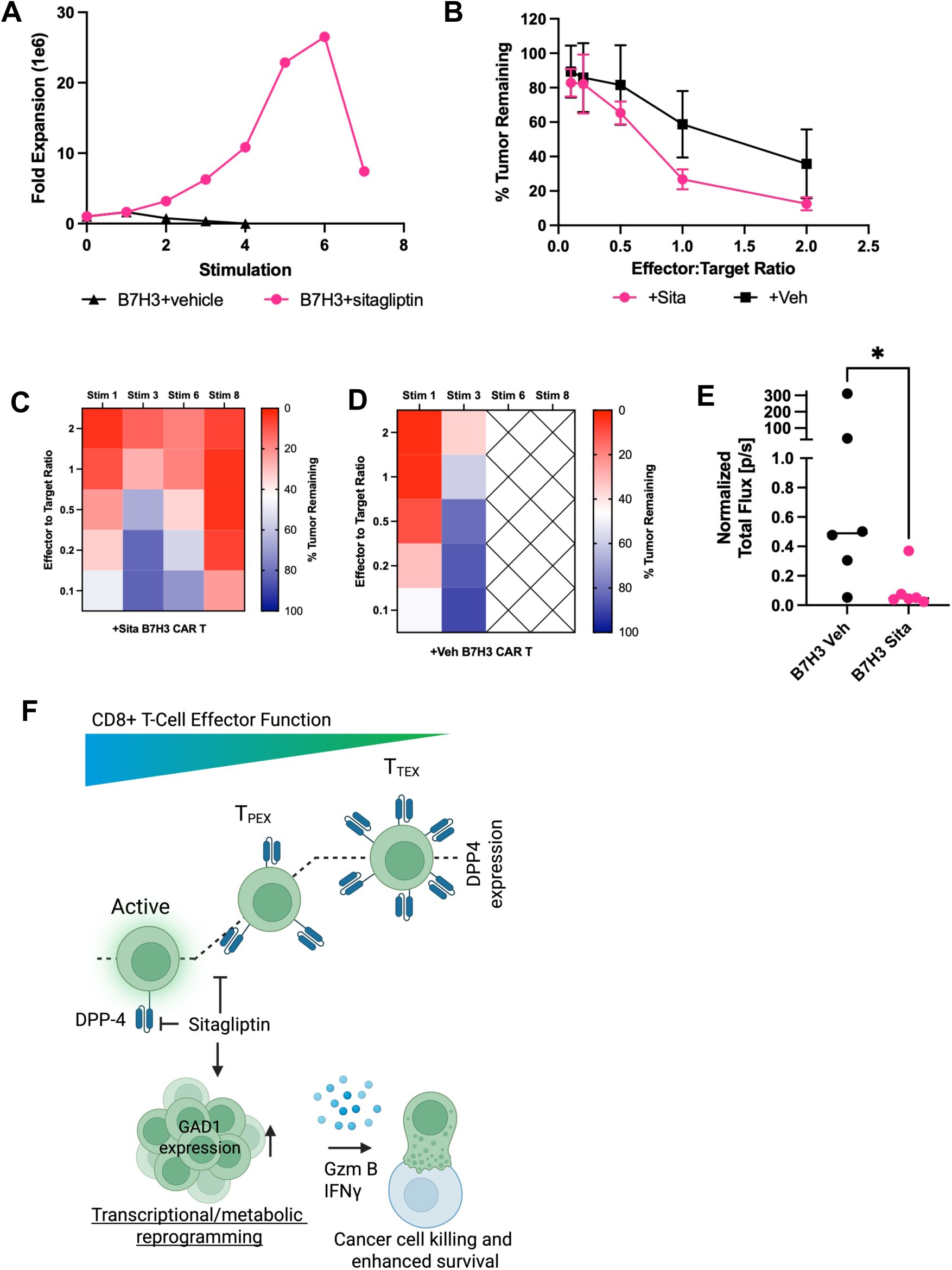
DPP4 inhibitors improve CAR-T Effector Function. Mouse CAR T-cells targeting B7-H3 were cultured with cancer cells expressing B7-H3 or empty vector at **(A)** 2:1 effector:target (E:T) ratio collected after 3-4 days, and co-cultured with fresh cancer cells until CAR T-cells stopped persisting in culture, or **(B)** at decreasing E:T ratios during the 3^rd^ stimulation to determine cancer cell cytotoxicity with MTS reagent after 72-hour co-culture. Heatmap demonstrating the killing capacity of either **(C)** sitagliptin- or **(D)** vehicle-treated B7-H3 CAR T-cells at reducing E:T ratio across different restimulations. **(E)** Normalized flux signal on day 35 post-intracranial injection n=4; * p<0.05 by t-test. **(F)** Diagram depicting the mechanism by which DPP-4 inhibition promotes metabolic/transcriptomic rewiring to enhance the effector function of CD8^+^ T cells.

## Discussion

Immunotherapies have not been uniformly successful across cancers with only a subset of patients experiencing long-term benefit even in immunotherapy-responsive histologies^71–73^. This is partly due to the prevalence of metabolically compromised CD8^+^ T cells and the inability of immune checkpoint inhibitors to reinvigorate T cell metabolism^16,17,74^. Similarly, exhaustion of adoptively transferred T cells, which limits their therapeutic efficacy, is accompanied by metabolic dysfunction^6–15^. Thus, identifying functional regulators of T cell metabolism holds potential for improved immunotherapy efficacy. In this study, we describe a noncanonical role for DPP-4 as a metabolic checkpoint molecule by demonstrating that pharmacological inhibition of DPP-4 improves GBM outcomes and sustains CD8^+^ T cell effector function.

Given T cell exhaustion is driven by mitochondrial dysfunction, metabolic reprogramming has been explored as a potential cancer immunotherapy strategy in preclinical models. While modulation of co-stimulatory signaling (e.g., CD28), key transcriptional factors and kinases (such as c-Myc, HIF1α, and mTOR), metabolic enzymes (including PGC-1α, LDH, and IRE1α/XBP1), or metabolite availability (e.g. L-arginine) has been effective^6–13,24,33–40^, the translational success has been limited by the predominantly intracellular localization of the targets, and the lack of safe and effective drug candidates. In contrast, DPP-4 is a membrane-bound and well-drugged protein. Based on our observation that DPP-4 is highly expressed by CD8^+^ T cells and its levels further increase with exhaustion, we interrogated the downstream effects of sitagliptin, an FDA-approved drug with a safe clinical profile and minimal drug-drug interactions. Our results indicated that DPP-4 inhibition metabolically reprograms CD8^+^ T cells. T cell metabolic signaling could be mediated by “top-down” TCR-dependent and “bottom-up” TCR-independent pathways^75^. We observed phosphorylation of the co-stimulatory molecule CD28, TCR downstream mediators, LCK and ZAP70, and the transcription factor NF-κB, as well as transcriptional activation of lymphocyte pathways pointing to a “top-down” regulation of T cell metabolism upon DPP-4 inhibition. In parallel, we also observed an upregulation of GAD1 with sitagliptin treatment. While the role of GAD1 in T cell metabolism w^13^as previously unknown, earlier studies demonstrated that the uptake of glutamine and its breakdown to glutamate is important for T cell activation and function^76,77^. Since GAD1 is involved in feeding glutamate into the TCA cycle, our results suggest that DPP-4 inhibition also initiates “bottom-up” metabolic signaling. The consequence of this broad reprogramming was enhanced metabolic fitness of T cells as evidenced by upregulation of both oxidative phosphorylation and glycolysis. Heightened metabolism can drive dysfunction via generation of harmful ^15^biproducts, but we observed that two essential cellular pathways that maintain redox homeostasis, pentose phosphate pathway and glutathione metabolism^11,78^, were enriched in our metabolomics dataset of sitagliptin vs vehicle-treated T cells. As such, we did not observe an increase in superoxide levels upon DPP-4 inhibition. Collectively, our results expand the role of DPP-4 from systemic to cellular metabolism.

While DPP-4 was initially identified as a regulator of incretin peptide hormones, there is a growing appreciation for the immunological role of DPP-4 with potential pro- or anti-tumorigenic consequences. Earlier studies demonstrated that high DPP-4 expression in a subset of CD4^+^ T cells is linked to increased anti-tumor immunity^79,80^ and that DPP-4 expression is also required for CD4^+^ T cell memory differentiation^81^. This immunostimulatory role of DPP-4 was in part mediated by its ability to sequestrate the adenosine deaminase (ADA) receptor and promote adenosine decomposition^45,82^. In our analysis of patient specimens and human donors, we observed that DPP-4 was variably expressed by GBM-infiltrating CD4^+^ T cells between the two datasets. On the other hand, we observed that CD8^+^ memory T cells had higher DPP-4 levels similarly to CD4^+^ memory T cells. Importantly, temporal DPP-4 inhibition promoted the differentiation of CAR T-cells to central memory phenotype, while delaying exhaustion of mouse CD8^+^ T cells. Collectively, these results indicate that inhibition of DPP-4 does not restrain memory formation and warrant studying its differential role in exhausted vs. memory T cells. While ligands, co-receptors, and direct binding partners of DPP-4, including ADA, in CD8^+^ T cells remain to be studied, the distinct role of DPP-4 in CD4^+^ vs CD8^+^ lymphocytes point to a cell-type-specific and contextual effect. In fact, it was recently reported that sitagliptin enhances dendritic cell antigen processing and subsequent T cell activation in colon adenocarcinoma models^83^, while also promoting tumor control through CCL11-mediated eosinophil infiltration in melanoma^84^. Our earlier results had implicated immunosuppressive myeloid cells as DPP-4 inhibitor targets in GBM^41^. In addition, sitagliptin was shown to decrease proliferation of CD4^+^ T_reg_ population^85,86^, and extend the bioactivity of CXCL10 to promote lymphocyte migration in ovarian cancer, pancreatic ductal adenocarcinoma, and hepatocellular carcinoma^48–50^. In addition to these defined roles, many cytokines, chemokines, and neuropeptides contain DPP-4 target sites, though functional implications of DPP-4 cleavage remain unclear. These observations support investigating common and divergent mechanisms of local and systemic immune regulation by DPP-4 across disease states. As such, conditioning CAR T cells ex vivo offers a therapeutic advantage by precisely regulating DPP-4 activity before infusion, thereby enhancing functional potency while reducing the risk of off-target effects.

While future studies will expand on how DPP-4 functions, our retrospective analysis of patient data indicated that gliptin use is associated with better GBM outcomes. Depletion experiments in preclinical models suggested that CD8^+^ T cells play a central role in this phenotype. Metformin, another anti-diabetic drug, has been reported to inhibit mitochondrial complex I, increase ADP, and activate AMPK in CD8^+^ T cells, enhancing persistence and accumulation of tumor-infiltrating lymphocytes and promoting IFNγ production^87–89^. Furthermore, the combination treatment of metformin and the mTOR inhibitor, rapamycin, promoted CAR T-cell function in preclinical models of GBM^89^. However, metformin use had no effect on GBM survival when compared to SOC alone. In contrast, investigations in colorectal or airway cancer patients suggest that gliptins associate with progression-free survival, while metformin reduces colon cancer overall mortality ^90,91,92^. Thus, given that sitagliptin and metformin both support T cell metabolism via distinct mechanisms, prospective clinical trials are necessary to discern their specific immune effects and potential as immunotherapeutic agents across different tumor types in combinatory studies.

In summary, our study highlights an unrecognized role of DPP-4 as a CD8^+^ T cell checkpoint molecule. Considering DPP-4 exists as a secreted exopeptidase, future studies can expand into the exploration of systemic DPP-4 levels/activity or its downstream targets as biomarkers of treatment response. Importantly, our results also provide a strong rationale for repurposing DPP-4 inhibitors in cancers, which are recognized for T cell dysfunction, as a single agent or in combination with established immunotherapies.

## Methods

### Cell lines and mice

The SB28 and SB28-OVA cell lines were obtained from Dr. Hideho Okada (University of California, San Francisco). These lines were maintained in RPMI 1640 (Corning #10-040-CV) supplemented with 10% FBS (Corning #4500-736) and 1% Antibiotic-Antimycotic (anti-anti, Gibco #15240062). Cell lines were routinely tested for mycoplasma with a Mycoplasma Test Kit (Lonza #LT07-118) and were not used beyond passage 25.

The U87MG cell line was purchased from the American Type Culture Collection (ATCC #HTB-14) and cultured in MEM (Richter’s modification) with pyruvate, GlutaMAX-1, HEPES, and penicillin/streptomycin (Thermo # 15140122) and supplemented with 10% fetal bovine serum (FBS). This cell line was engineered to express EGFRvIII protein, green fluorescent protein (GFP), and click beetle green (CBG) luciferase protein through single-cell purification and expansion.

Four-week-old C57BL/6 male and female mice (JAX Stock #000664) were purchased from the Jackson Laboratory as needed. P14 TCR transgenic mice expressing a TCR specific for the LCMV Dbgp33-41 epitope were acquired from Dr. Erietta Stelekati (University of Miami) and bred in house. C57BL/6-Tg(TcraTcrb)1100Mjb/J (OT-I, JAX Stock #003831) mice were purchased from Jackson Laboratory and bred in house. All animals were housed in the DVR facility at University of Miami Miller School of Medicine.

### In vivo tumor studies

C57BL/6 mice were intracranially injected at 4-6-weeks old with 20,000 SB28 in 10 μL RPMI media into the left cerebral hemisphere 2 mm caudal to the coronal suture, 3 mm lateral to the sagittal suture at a 90° angle with the murine skull to a depth of 2.5 mm, using a stereotaxis apparatus (Kopf). Mice were then subcutaneously injected with 3.5 mg/kg buprenorphine for analgesic purposes. Mice were monitored for neurological symptoms, including lethargy, hunched posture, loss of balance, and head tilt that would indicate tumor burden and determine endpoint.

For sitagliptin (Cayman Chemical #13252) treatment, SB28-bearing mice were provided with 4 g/L sitagliptin-containing water starting at the day of tumor implantation. For CD8^+^ T cell depletion, mice were intraperitoneally injected with 200 μg of InVivo Mab anti-mouse CD8a (BioXcell, clone: 2.43) or isotype control (BioXcell, Clone: LTF-2) one day before intracranial injection and once again on the day of the procedure. Mice were then intraperitoneally injected every 5 days with 100 μg antibody until endpoint.

### LCMV infection model

Wild-type C57BL/6 (CD45.2⁺) recipient mice were infected with 2 x 10^5^ plaque forming units (PFU) of LCMV Armstrong intraperitoneally (i.p.) or with 4 x 10^6^ LCMV clone 13 intravenously (i.v.). 1 × 10⁴ naïve P14 TCR-transgenic CD8⁺ T cells (CD45.1⁺) were adoptively transferred intravenously into recipient mice one day prior to infection (day –1). Body weights were monitored twice weekly post-infection, and mice were euthanized if they experienced greater than 30% weight loss. Peripheral blood samples were collected on days 16 and 21 post-infection to assess expansion of donor P14 cells. Serum was isolated for viral titer analysis by Quantitative real-time PCR to test LCMV glycoprotein (GP1)-specific mRNA.

For sitagliptin (Cayman Chemical #13252) treatment in chronic LCMV Model, wild-type C57BL/6 (CD45.2⁺) recipient mice were infected with 4 x 10^6^ LCMV clone 13 intravenously (i.v.) on day 0.1 x 10^5^ naïve P14 cells were transferred at day -1. Mice were treated with 20 mg/kg sitagliptin intraperitoneally for two rounds of 5 consecutive days beginning day 3 post-infection, followed by a 2 day off period in between the two rounds. Spleens were harvested 24 hours following the final dose and processed as described below for downstream immunophenotyping. Mouse body weights were monitored daily during treatment. Peripheral blood was collected on day 7 (during treatment) and at terminal time points.

### Immunophenotyping of mouse splenocytes and tumors

Spleens were harvested from either healthy 6-week-old C57BL/6 mice or on day 14^th^ after intracranial injection for tumor bearing mice, and as specified above for LCMV models. Splenocytes were then isolated using a 40 μm cell strainer. Cells were span at 340 *g*, resuspended in 500 μL PBS, and aliquoted for flow staining. When applicable, tumors were also extracted and enzymatically digested with Collagenase Type IV, 1 mg/mL (Stem Cell Technologies #7900) containing DNase I diluted to 1:1000 (Thermo #18047019) at 37°C for 15 mins, with intermittent shaking every 5 minutes. Following incubation, tumors were mechanically dissociated by passage through a 70 μm strainer to create a single cell suspension. Strainers were washed with PBS and cells were span at 340 *g* for 5 minutes. Pellets were treated with Red Blood Cell (RBC) lysis buffer (Biolegend #420302) for 5 minutes at room temperature, span again, and then resuspended in 500 μL PBS and aliquoted for flow staining. After processing, all cells were stained with Zombie Aqua (Biolegend #423101) at 1:500 in PBS for 10 minutes to distinguish live populations and then incubated in FcR mouse blocking buffer (Miltenyi Biotec #130-092-575) at 1:50 in FACS buffer (PBS supplemented with 2%BSA) for 5 minutes. Antibody containing FACS buffer was then added for extracellular marker staining, and cells were incubated on ice in the dark for 20 minutes. Immune populations were defined as follows: Dendritic cells (DCs; CD45^+^, CD11c^+^), macrophages (CD45^+^, CD11b^+^, CD68^+^, F4/80^+^), granulocytic myeloid-derived suppressor cells (gMDSCs; CD45^+^, CD11b^+^, Ly6C^low^, Ly6G^+^), monocytic myeloid-derived suppressor (mMDSCs; CD45^+^, CD11b^+^, Ly6G^-^, Ly6C^high^), monocytes, (CD45^+^, CD11b^+^, Ly6G^-^, Ly6C^high^, IA/IE^+^), B cells (CD45^+^, B220^+^), CD4^+^ T cells (CD45^+^, CD3^+^, CD4^+^), and CD8^+^ T cells (CD45^+^, CD3^+^, CD8^+^). For LCMV models, DPP4 expression levels were quantified by flow cytometry in the following CD8^+^ T cell populations from LCMV-Arm infected mice: Naive (CD44⁻CD62L⁺), activated (CD44⁺CD62L^-^), effector (CD127^-^KLRG1^+^), and memory (CD127⁺KLRG1⁻). In LCMV-Clone13 infected mice, exhausted T cells were identified as CD69⁺Ly108^-^ P14 cells. All antibodies were used at 1:100 unless otherwise specified (**Table S1**).

### Human PBMCs processing and immunophenotyping

LRS (Leukocyte Reduction System) chamber were purchased from Suncoast Community Blood Bank, Inc. Contents were extracted by flushing chambers with PBS without Ca2^+^ or Mg2^+^. Blood was diluted to 45 mL and then 15 mL layered on top of 15 mL of Ficoll Paque Plus (Sigma, #GE17-1440-02, density: 1.077 g/mL). Samples were span at 2200 rpm with normal acceleration, and zero break for 20 minutes at room temperature. After spin, buffy coat PBMCs were carefully removed with a transfer pipette and span for 5 minutes at 1500rpm. Pellet was then treated with Red Blood Cell (RBC) lysis buffer (Biolegend #420302) for 5 minutes at room temperature, and then repetitively washed 3 times and filtered through a 40 μm cell strainer. PBMCs were then either frozen for downstream functional assays or freshly immunophenotyped as follows: at least 2 million cells were stained with Zombie Aqua at 1:500 in PBS for 10 minutes to distinguish live populations and then incubated in FcR human blocking buffer (Miltenyi Biotec #130-059-901) at 1:50 for 5 minutes. Antibody containing FACS buffer (PBS supplemented with 2%BSA) was then added for extracellular marker staining, and cells were incubated on ice in the dark for 20 minutes. CD8^+^ T cell populations were defined as: CD45^+^, CD3^+^, CD8^+^ plus effector (CD45RA^+,^ CD62L^-^), central memory (CD45RA^-^, CD62L^+^), effector memory (CD45RA^-^, CD62L^+^), and stem-like memory (CD45RA^+^, CD62L^+^, CD95^+^).

### T cell phenotyping from high-grade gliomas

We collected surgical specimens from 4 patients with clinical diagnosis of high-grade glioma. Specimens were minced and then enzymatically digested with Collagenase Type IV, 1 mg/mL (Stem Cell Technologies #7900) containing DNase I diluted to 1:1000 (Thermo #18047019) at 37°C for 45 mins, with intermittent shaking every 5 minutes. Following incubation, tumors were mechanically dissociated by passage through a 70 μm strainer to create a single cell suspension. Strainers were washed with PBS and cells were span at 340 *g* for 5 minutes. Pellets were treated with RBC lysis buffer (Biolegend #420302) for 5 minutes at room temperature, span again, and then stained with Zombie Aqua (Biolegend #423101) at 1:500 in PBS for 10 minutes. After live/dead stain, cells were incubated in FcR human blocking buffer (Miltenyi Biotec #130-059-901) at 1:50 for 5 minutes. Antibody containing FACS buffer (PBS supplemented with 2%BSA) was then added for extracellular marker staining. Cells were fixed and permeabilized using True-Nuclear Transcription Factor Buffer Set (Biolegend #424401). In brief, fixing reagent was diluted 1:3 with fix diluent buffer and added to cells for 30 minutes on ice. Cells were then washed with FACS buffer and resuspended in 1X permeabilization buffer diluted in MQ H_2_O. After permeabilization, cells were resuspended in antibody containing permeabilization buffer and incubated in the dark on ice for 60 minutes. CD8^+^ T cells were defined as CD45^+^, CD3^+^, CD8^+^ plus TIGIT^+^, PD1^+^, and TOX^+^ as exhaustion markers.

### Cancer Cell Apoptosis Assay

50,000 cancer cells were seeded in 96-well round bottom plates and treated with either vehicle control or 100μM sitagliptin. Cells were incubated for 48 hours and then stained at room temperature in the dark with 1:250 viability stain 7-AAD (Biolegend #420404) in Annexin Binding Buffer (Biolegend #79998) for 10 minutes. Cells were washed and resuspended in 150 μL PBS for analysis on a CytoFLEX LX.

### CD8^+^ T cell functional assays

#### Proliferation assay

Mouse CD8^+^ T-cells were isolated from freshly harvested spleens of C57BL/6 mice using CD8a (Ly-2) MicroBeads (Miltenyi #130-114-044), following manufacturer’s instructions. After isolation, T cells were stained with 10 μM 5-(and 6)-Carboxyfluorescein diacetate succinimidyl ester (CFSE, Biolegend #423801) in the dark for 5 minutes at 37°C and reaction was quenched with ice-cold RPMI containing 10% FBS. Cells were then seeded at 1 x 10^5^ cells/well in a 96-well round bottom plate and activated in T-Cell media (RPMI + 10% FBS + 1x anti-anti) containing 100 IU/mL mouse IL-2 (Biolegend #575402), Dynabeads™ Mouse T-Activator CD3/CD28 for T-Cell Expansion and Activation (Thermo #11453D), and 100 μM sitagliptin, 100 μM L-Allylglycine (Cayman Chemical #23348), combination treatment, or vehicle control (PBS) for either 4 or 6 days. CFSE dilution was determined using a CytoFlex flow cytometer. For human specimens, PBMCs were acquired as previously described. Frozen stocks were then thawed, and CD8^+^ T cells were isolated using human CD8 MicroBeads (Miltenyi #130-045-201). Cells were either stained with 10 μM CFSE as done for mouse cells or manually counted on day 6 post-isolation. Human CD8^+^ T cells were expanded in human T cell media (RPMI + 5% human AB serum, Sigma #H4522, + 1x Penicillin-Streptomycin, Corning #30-002-Cl) containing 100 IU/mL human IL2 (Biolegend #589102), Dynabeads™ Human T-Activator CD3/CD28 for T-Cell Expansion and Activation (Thermo #11161D), and 100 μM sitagliptin, 100 μM L-Allylglycine, combination treatment, or vehicle control (PBS).

#### Cancer cell killing assay

CD8^+^ T cells were MACS-sorted from OT-I splenocytes. T cells were then seeded in 12-well plates at 1.5 x1 0^6^ per well and activated for 2 days with 5 ng/mL IL-7 (PeproTech #200-07-10UG), 5 ng/mL IL-15 (PeproTech #210-15-10UG), and 10 ng/mL OVA peptide 257-264, (InvivoGen #vac-sin) in the presence of either 100 μM sitagliptin or vehicle control. After activation, T cells were co-cultured with either OVA-expressing SB28 (GFP^+^) or control SB28 (GFP^+^) at 1:1, 1:5, and 1:10 E:T (effector:target) ratios in 96-well round bottom plates. The number of resuspended cells added was adjusted to seed a total of 120,000 cells per well. After 6 hours, cells were stained at room temperature in the dark with 1:250 viability stain 7-AAD (Biolegend #420404) in Annexin Binding Buffer (Biolegend #79998) for 10 minutes. Cells were washed and stained on ice with anti-mouse CD8 at 1:100 in FACS buffer (PBS, 1%BSA) for 20 minutes. Cells were washed and resuspended in 150 μL PBS for analysis on a CytoFLEX LX.

#### Cytotoxic mediator production

CD8^+^ T cells were isolated from OT-I splenocytes and seeded in round-bottom 96-well plates at 1 x 10^5^ per well. Cells were activated with IL-7, IL-15 and OVA peptide as above in the presence of either 100 μM sitagliptin or vehicle control and media was replaced every day for five days. On day 6, cells were treated with GolgiPlug (BD #555029, 1:1000 dilution) and GolgiStop (BD #554724, 1:1500) in RPMI (10% FBS and 1x anti-anti) and incubated for 4-6 hours at 37°C with 5% CO_2_. After incubation, cells were extracellularly stained as previously described and then fixed and permeabilized using True-Nuclear Transcription Factor Buffer Set (Biolegend #424401) as described in the manufacturer’s protocol. In brief, fixing reagent was diluted 1:3 with fix diluent buffer and added to cells for 30 minutes on ice. Cells were then washed with FACS buffer and resuspended in 1X permeabilization buffer diluted in MQ H_2_O. After permeabilization, cells were resuspended in 1:100 anti-IFN and anti-Granzyme B antibody containing permeabilization buffer and incubated in the dark on ice for 60 minutes.

#### Seahorse Assay

After MACS isolation, murine CD8^+^ T-Cells were seeded at 1 x 10^6^ cell/well in a 12-well plate and activated in T-Cell media (RPMI+10%FBS+1X anti-anti) containing 100 IU/mL mouse IL-2 plus anti-CD3/CD28 Dynabeads and 100 μM sitagliptin, 100 μM L-Allylglycine, combination treatment, or vehicle control (PBS) for 4 days. For human specimens, PBMCs were acquired as previously described. Cells were expanded in human T-Cell media, containing 100 IU/mL human IL-2, plus anti-CD3/CD28 Dynabeads, and 100 μM sitagliptin, 100 μM L-Allylglycine, combination treatment, or vehicle control (PBS) for 4 Days. On day of assay, cells were collected, counted, and seeded at 100,000-150,000 per well in seahorse plates (Agilent #103794-100) previously coated with 3.5 μg/cm^2^ Cell Tak solution (Corning #354240). Cells were then subjected to the T-Cell Fitness Assay (Oligomycin A, Selleckchen #S1468, at 1.5μM, Bam15, Sigma #SML1760, at 2.5μM, and Rotenone/antimycin A, Sigma #557368 and #A8674, at 1μM) on Agilent Seahorse XF Pro Analyzer.

#### TMRM, MitoSOX, and 2-NBDG staining

After isolation cells were seeded at 1x10^5^ cell/well in a 96-well round bottom plate and activated as previously described. After treatment, media was replaced with full T-Cell media containing and treated as follows: (1) 100 nM TMRM Reagent (Thermo #I34361), 30 minute incubation, (2) 500 nM MitoSOX Red (Thermo #M36008), 60 minute incubation, or (3) 50 μM 2-NBDG (2-(N-(7-Nitrobenz-2-oxa-1,3-diazol-4-yl)Amino)-2-Deoxyglucose, Thermo #N13195) for 30 minutes. All incubations were performed at 37°C with 5% CO_2_.

### Characterization of sitagliptin-mediated global changes

#### Bulk RNA sequencing

After MACS isolation, mouse CD8^+^ T-cells were seeded at 1 x 10^6^ cell/well in a 6-well plate and activated in full T-Cell media containing 100 IU/mL mouse IL2, anti-CD3/CD28 Dynabeads and 100 μM sitagliptin or vehicle control (PBS) for 4 days. On Day 4, cells were collected and RNA was isolated using RNeasy Kit (Qiagen #74104). Total RNA was sent to Novogene for mRNA sequencing using Illumina sequencing platform. Top differentially up-regulated pathways in sitagliptin-treated CD8^+^ T cells based on genes expressed >2-fold with p<0.05.

#### Phosphoproteomic antibody array

Mouse CD8^+^ T-cells were isolated and activated as described above for 18 hours. After overnight activation and treatment, cells were collected, span at 340 *g* for 5 minutes, washed with PBS, and then flash frozen. Cell pellets containing at least 5 x 10^6^ cells per samples were sent to Fullmoon BioSystems to perform T-Cell Receptor Phospho Antibody Array (FullMoon BioSystems #PTC188). In brief, cell pellets were lysed with beads and extraction buffer for 60 minutes with repeat vortexing for 30 seconds at 10-minute intervals. Once supernatant is isolated, proteins are labeled with 100 uL of DMF to 1 mg of Biotin Reagent to give a final concentration of 10 μg/μL. Coupling Chamber containing antibody microarray is now exposed to labeled protein samples and incubated on an orbital shaker for 2 hours at room temperature. After washing, Cy3-Streptavidin, 1 mg/ml, is used to detect bound proteins labelled with biotin. Slides are dried with compressed nitrogen and imaged on an Axon GenePix Array Scanner. Protein targets that were upregulated or phosphorylated >1.2-fold were functionally annotated with https://maayanlab.cloud/Enrichr/.

#### Metabolomic analysis

Mouse CD8^+^ T-cells were isolated and then seeded at 1 x 10^5^ cell/well in a 96-well round bottom plate and activated in full T-Cell media containing 100 IU/mL mouse IL2, Dynabeads™ Mouse T-Activator CD3/CD28 for T-Cell Expansion and Activation, and 100 μM sitagliptin or vehicle control (PBS) for 2 days. After treatment, cells were quickly collected by centrifugation at 340 *g* for 5 minutes at 4°C and supernatant removed. Cell pellets were placed on dry ice and 1 mL of LCMS grade 80%MeOH/H20 (precooled at -80°C for at least 1 hour) was added. Samples were then incubated at -80°C for 3-4 hours. After incubations, samples were placed on ice and vortexed several times to achieve full extraction. Samples were then span at 20,000 *g* for 10 minutes at 4°C and supernatants transferred to a new tube. Metabolite containing supernatant was speed-vac dried overnight and then stored at -80°C for downstream processing. Samples were reconstituted in 50 µL of LCMS grade water for reverse-phase chromatography and in 80% LCMS grade acetonitrile for HILIC-based chromatography.

Agilent 1290 ultrahigh-performance liquid chromatography (UHPLC) and 6495C QqQ mass spectrometer were employed to separate and detect metabolites. Agilent Masshunter acquisition software 12.1 was used to acquire data. The following method parameters were used for reverse phase chromatography: 2 μL injection volume, 0.250 mL/Min flow, UHPLC Guard (P.N.821725-907) and Agilent Zobrax extend C18 column (P.N. 759700902), 35.0 °C column temperature, Solvent A (LCMS grade water with 3% LCMS grade methanol, 10mM Tributylamine (TBA), and 15mM glacial acetic acid) and solvent C (LCMS grade methanol with 10mM Tributylamine (TBA), and 15mM glacial acetic acid) for chromatographic separation, and solvent D (100% LCMS grade acetonitrile) for column wash. The following gradient was used: 100% A with 0.250 mL/Min flow at 0 to 2.5 min, 80% A and 20% C with 0.250 mL/Min flow at 7.5 min, 55% A and 45% C with 0.250 mL/Min flow at 13 min, 1% A and 99% C with 0.250 mL/Min flow at 20 min, 1% A and 99% C with 0.250 mL/Min flow at 24 min, 1% A and 99% D with 0.250 mL/Min flow at 24.05 min, 1% A and 99% D with 0.250 mL/Min flow at 27 min, 1% A and 99% D with 0.800 mL/Min flow at 27.5 min, 1% A and 99% D with 0.800 mL/Min flow at 31.35 min, 1% A and 99% with 0.600 mL/Min flow D at 31.5 min,100 % A with 0.400 mL/Min flow at 32.25 min, 100 % A with 0.400 mL/Min flow at 39.9 min, 100 % A with 0.250 mL/Min flow at 40 min. The QqQ was run in dynamic multiple reaction mode (dMRM) with following parameters: negative electrospray ionization, 600 ms cycle time, 150 °C gas temperature with 13 L/min flow, Nebulizer at 45 psi, 325 °C of Sheath Gas Temperature with 12 L/min flow, 2000 V of capillary and 500 V of Nozzle voltage.

The following method parameters were used for HILIC chromatography: 2 μL injection volume, 0.400 mL/Min flow, Agilent poroshell 120 HILIC-Z 2.7 µm, 2.1 x 150mm column (P.N. 683775-924), 15.0 °C column temperature, Solvent A (LCMS grade water with 20 mM ammonium acetate) and solvent B (100% LCMS grade acetonitrile) for chromatographic separation. The following gradient was used: 10% A and 90 % B with 0.400 mL/Min flow at 0 to 1 min, 22% A and 78 % B with 0.400 mL/Min flow at 8 min, 40% A and 60 % B with 0.400 mL/Min flow at 12 min, 90% A and 10 % B with 0.400 mL/Min flow at 15 min, 10% A and 90 % B with 0.400 mL/Min flow at 18 min, 10% A and 90 % B with 0.400 mL/Min flow at 19 min, 10% A and 90 % B with 0.500 mL/Min flow at 19.1 min, 10% A and 90 % B with 0.500 mL/Min flow at 22 min, 10% A and 90 % B with 0.400 mL/Min flow at 22.1 min, 10% A and 90 % B with 0.400 mL/Min flow at 23 min. The QqQ was run in dynamic multiple reaction mode (dMRM) with following parameters: positive electrospray ionization, 500 ms cycle time, 200 °C gas temperature with 14 L/min flow, Nebulizer at 50 psi, 375 °C of Sheath Gas Temperature with 12 L/min flow, 3000 V of capillary voltage. Relative abundance of each metabolite was quantified by using Agilent Quantitative software 12.1.

### Human CAR T-cell studies

#### Lentiviral vector production and T cell transduction

Lentiviruses were packaged in HEK293T cells using a split genome approach and tittered using SUPT1 cells (ATCC #CRL-1942). Normal human T cells were obtained from the Human Immunology Core at the University of Pennsylvania, using de-identified healthy donors under an IRB-approved protocol and were transduced using lentiviral vectors. T cells were stimulated for 5 days with Dynabeads Human T-Activator CD3/CD28 (Life Technologies at a bead-to-cell ratio of 3:1. After 24 hours of stimulation, T cells were transduced with 2173 41BBz lentivirus with a multiplicity of infection (MOI) of 5. The T cell concentration was determined using a Coulter Multisizer (Beckman Coulter) and maintained at 0.7 x 10^6^ cells/mL until fully rested at approximately 300 fl in volume. The cells were cultured in R10 media (RPMI-1640 with GlutaMAX-1, pyruvate, HEPES, and penicillin/streptomycin supplemented with 10% FBS) with 30 IU/mL recombinant human IL-2. The CAR T cells were then cryopreserved in a mixture of 90% FBS and 10% DMSO for future use.

#### Impedance cytotoxicity assays

U87vIII target tumor cells were seeded at 5 x 10^4^ into Axion Biosystems microelectrode containing 96-well plates (Axion Biosystems). The impedance plate was prepared prior to experimentation by coating with 0.1 mg/mL poly-D lysine (Gibco # A3890401) overnight at 37℃. After coating was complete, wells were washed with diH2O and overlaid with 100 μL of the culture medium. The plate was then placed on the Axion Biosystems ZHT analyzer to establish a baseline reading of the background impedance without cells present. After baseline was obtained, the plate was removed from the analyzer and was seeded with target cells in a volume of 150 μL/well. After cell plating, the plate was left in the cell culture hood for 1 hour at room temperature to ensure settling and attachment of the cells down to the microelectrodes on the bottom surface. The plate was then returned to the analyzer and data collection began. Data were collected every 1 minute for 24 hours for cell monolayer growth measurement. For cytotoxicity assessment, 2173 41BBz CAR T cells and UTD CAR T cells were added with and without sitagliptin or media only was added for a total volume of 200 μL/well. Sitagliptin was added at a concentration of 100 μM. Changes in impedance are reported as the resistive component of the complex impedance. Using AxIS Z software (Axion Biosystems), all data are corrected for “media alone” to remove small changes in media only impedance over time and then normalized to the impedance at the time of addition of effector cells. The percent cytolysis calculations utilize the no treatment control and full lysis controls to determine percent of target cell cytolysis as follows:

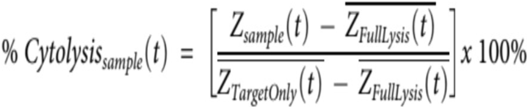

### Mouse CAR T-Cell studies

#### Generation

Spleens were collected from 6- to 8-week-old C57BL/6 mice and passed through a 70 μm cell strainer using the back of a syringe plunger. Red blood cells were then lysed using ACK (ammonium-chloride-potassium) lysis buffer (Life Technologies #A10492-01), washed, and pelleted. CD3^+^ murine T cells were then collected using the Pan T-Cell Isolation Kit II (Miltenyi Biotec #130-095-130) as per the manufacturer’s protocol. Isolated T cells were washed and activated using plate-bound CD3/CD28 antibodies (BD Biosciences #553057 and #553294 respectively) in complete RPMI media supplemented with 50 IU/mL of human IL-2 (Peprotech #200-02). T cells were activated in the presence of 100 μM of sitagliptin or vehicle control.

On Day 2 post-activation, T cells were transduced with retroviral particles encoding a mB7H3 chimeric antigen receptor (CAR) construct^93^ using RetroNectin-coated plates (Clontech #50-444-033) in complete RPMI media supplemented with 50 IU/mL of IL-2 and 100 μM of sitagliptin or vehicle. Non-transduced (NT) T cells were prepared by adding activated T cells on RetroNectin-coated wells without retroviral particles. After 2 days, CAR T cells were harvested and expanded with complete RPMI media supplemented with 50 IU/mL of IL-2 and 100 μM of sitagliptin or vehicle control. CAR detection was performed on Day 5 post-transduction using flow cytometry. Experiments were performed on Days 5-6 post-transduction. T cells are supplemented with sitagliptin 3 times prior to use in functional assays.

#### CAR T-cell phenotyping and transduction efficiency

The B7-H3-CAR was detected on T-cells with FITC-anti-human IgG, F(abʹ)2 fragment-specific antibody (Jackson ImmunoResearch #109-096-006). CAR positive cells were stained for phenotyping and exhaustion marker analysis at five days post-transduction.

#### CAR T-cell Stimulation Assay

For repeat stimulation assays, 5 x 10^5^ (GL261) tumor cells were plated in a 24-well tissue culture plate and allowed to adhere for 2-4 hours at 37°C. T-cell cultures (sitagliptin-treated and vehicle-treated CAR T-cells) were washed once with cytokine-free media before addition to assay plate. A total of 1 x 10^6^ or 5 x 10^5^ T-cells were then added to the wells containing tumor cells at 2:1 E:T ratio. After 3-4 days post-stimulation, T-cells were collected by gentle pipetting and quantified using a hemocytometer. Repeated stimulations were performed by re-plating fresh tumor cells and re-establishing co-culture with collected T-cells at the same E:T ratio every 3-4 days. Stimulations continued until T-cell proliferation and killing activity were no longer observed.

#### CAR T-cell cytotoxicity assay

Colorimetric MTS reagent [3-(4,5-dimethylthiazol-2-yl)-5-(3-carboxymethoxyphenyl)-2-(4-sulfophenyl)-2H-tetrazolium] (Promega #G3582) was used to quantify the cytotoxicity of CAR T-cells. Briefly, GL261 or SB28.B7H3.flag tumor cells were added to a 96-well tissue culture-treated plate and allowed to adhere for 2-6 hours. T-cell cultures post-manufacturing or post-stimulation were added to the tumor cells at varying E:T ratios (2:1, 1:1, 0.5:1, 0.25:1, and 0.125:1, performed by serial dilution). Wells containing media or tumor cells only were used as controls to determine the absorbance corresponding to background and no T-cell killing, respectively. T-cells and tumor cells were incubated for 72 hours at 37°C. All conditions were performed in triplicate. After 72 hours, T-cells were collected by gently pipetting up and down to avoid disrupting adherent tumor cells. MTS reagent was then diluted in complete media and added to each well, incubated at 37°C for 2 hours and absorbance measured at 492 nm by using a SPARK Plate Reader (Tecan). The percentage of live tumor cells was calculated using the following formula: [(Absorbance of sample- Absorbance of media only)/ (Absorbance of tumor only - Absorbance of media only)] x100.

#### Immunocompetent glioma model

Albino C57BL/6 mice (B6(Cg)-*Tyr^c-2J^*/J, 000058, The Jackson Laboratory) were purchased at 5 weeks of age. All animals used for tumor implantation were 6-week-old males. For intracranial implantation, mice were anesthetized and placed in a stereotactic rodent surgery platform. A 1-mm burr hole was drilled into the skull at 0.8 mm posterior and 1 mm to the right of the lambda. A total of 1x10^5^ KAPP.ffLuc cells in 1 μL of media were injected at 5.5 mm deep over 2 minutes as described in^ref^. Wound clips were then used to close the surgical site. At Day 10 post tumor implantation, animals were treated with 1x10^6^ effector cells suspended in 1 μL of RPMI via intra-cranial injection using the same burr hole coordinates. Mice were euthanized when they reached an endpoint defined by significant weight loss, signs of distress, or when recommended by the Children’s National Hospital veterinary staff. Each experiment involved 7 mice per group.

#### Bioluminescence imaging

Mice were injected intraperitoneally with AkaLumine hydrochloride (MedChemExpress, Monmouth Junction, NJ) (5 mM in PBS; 100 µL per mouse) 5 minutes before imaging, anesthetized with isoflurane (2.5% delivered in 100% O_2_ at 0.5 L/min), and imaged with a Xenogen IVIS-50 imaging system (Xenogen, Alameda, CA). The photons emitted from the luciferase-expressing cells were quantified using Living Image software (Caliper Life Sciences, Waltham, MA). Mice were imaged 4 and 7 days post-implantation and then weekly to track tumor burden.

### Retrospective GBM patient study

#### Summary of Methods

Real world data was obtained from linked Surveillance, Epidemiology, and End Results (SEER) data and Medicare claims for GBM cases diagnosed from January 1, 2008 to December 31, 2019 with corresponding Medicare claims through December 31, 2021. Analysis was limited to those that received surgical resection, radiation, and temozolomide (TMZ), leaving 2,543 individuals for analysis. Cox proportional hazard models were adjusted for sex, age, race/ethnicity, Charlson comorbidity score, and extent of resection (subtotal resection [STR], gross total resection [GTR], or unknown extent of surgery) to estimate hazard ratios (HR), 95% confidence intervals (95% CI), and p-values. Statistical significance was assessed at p<0.05. Survival was assessed comparing Metformin+TMZ, DPP4i+TMZ, and combination to TMZ only. In addition, survival comparing Metformin+TMZ to DPP4i+TMZ was assessed (**Fig. S1**).

#### Limitations

While Medicare data is a powerful resource for research, there are several important limitations. All individuals included in this analysis are ≥65 years, meaning that younger individuals are excluded. The median age of first diagnosis with GBM is ∼663, and as a result this should capture >50% of the population with GBM. While the vast majority of those ≥65 in the US are entitled to Medicare, there are several factors that affect whether claims are reported. Individuals in the workforce may retain employer-sponsored insurance as primary insurer and Medicare as secondary resulting in some claims being billed to the primary insurance. These data include only traditional Medicare, while many individuals select Medicare Advantage (MA) coverage through private insurance companies, which is paid for by Medicare but does not result in claims directly to CMS. While there are no substantial differences between those choosing traditional Medicare as opposed to MA,^94^ there may be significant differences between those with private insurance, MA, and those with traditional Medicare that would limit the generalizability of our findings. While SEER-Medicare contains nearly all claims billed through Medicare, previous research has shown that the information contained in the claims themselves is not always reliable or valid.^95–98^ Threats to the validity of claims may include under/over-estimates prescribed, patient compliance with billed treatments, and inconsistent use of billing codes, among others. A major limitation of any analysis using insurance claims data is the inability to identify conditions or medications which do not have an associated billing code, such as participation in a clinical trial or imaging evidence of disease progression. As a result, we are not able to assess potential interactions with unapproved treatments such as immunotherapies. Finally, the observational nature of our study precludes establishing causality between Metformin and DPP4i usage and improved GBM survival.

#### Extended Methods

The National Cancer Institute’s SEER program includes specially funded population-based cancer registries that collect data on newly diagnosed cancers within catchment area. For individuals that are enrolled in Medicare, existing SEER case data can be linked to Medicare claims. This linked dataset includes 15 SEER registries covering ∼24% of the US population^99^.

GBM cases were identified using International Classification of Diseases, Oncology 3rd edition [ICD-O-3] morphology codes 9440-9442/3, and ICD-O-3 topography codes C71.0-C71.9^100^. Individuals were excluded if: 1) qualified for Medicare for a reason other than age, including End-Stage Renal Disease and/or disability; 2) enrolled in a Health Maintenance Organization during the period of six months prior to twelve months after primary diagnosis with GBM; 3) not continuously enrolled in Medicare Parts A, B, and D for the period of 6 months prior to diagnosis through the end of follow-up; 4) missing race, sex, or follow-up time.

Primary site was categorized using ICD-O-3 morphology codes as supratentorial for primary sites C71.0-C71.4, infratentorial at primary sites C71.6-C71.7, and other for all other primary sites. Comorbidity score was estimated using the rule-out option of the SEER-Medicare SAS comorbidity macro (2021 version)^101^ using both International Classification of Disease version 9 or version 10 (ICD-9, ICD-10) diagnosis codes. Comorbidity scores were calculated with the Charlson weight, and categorized as a score of 0-1, 2 or 3. Cancer treatment patterns extent of surgical resection, radiation, chemotherapy (temozolomide) were identified using SEER data and claims from the MEDPAR, NCH, OUTPAT, Part D Event, and Durable Medical Equipment files and treatments were only included if they occurred within 3 months of initial diagnosis. Extent of resection was identified using SEER-provided variable, RX_SUMM_SURG_PRIM_SITE, and claims with Common Procedural Terminology (CPT) and ICD-9 and ICD-10 procedure codes.

In SEER, having a surgical procedure is based on the variable RX_SUMM_SURG_PRIM_SITE_1998 or the presence of a Medicare claim. Extent of resection is further classified into no surgery, gross total resection (GTR), subtotal resection (STR), unknown surgery, and other surgery. Using the SEER variable, extent of resection was identified as no surgery if they had a value of 0, GTR if they had a value of 30 or 55, STR if they had a value of 20-22 or 40, other surgery if they had a value of 10 or 90, and unknown surgery if they had a value of 99. When the SEER surgery variable is either 0 or 99, we further classified surgery type using the linked Medicare claims data. In the claims data, we are able to identify “biopsy only” claims based on ICD-9 or ICD-10 procedure code or CPT. For all other neurosurgical resection ICD-9 or ICD-10 procedure code or CPT, those were classified as other surgery, given we cannot distinguish from the procedure codes the extent of surgical resection.

Radiation therapy and chemotherapy were defined by ≥2 claims for the same code separated by at least one week started after the diagnosis date. Radiation therapy was identified using the SEER-provided radiation variable (RADIATION), with a value of ‘0’ or ‘7’ corresponding to no radiation, values 1-6 corresponding to some radiation, and all other values corresponding to missing data. When SEER radiation data were missing, the presence of radiation therapy in the linked Medicare claims data were used to identify radiation therapy. Temozolomide use was identified using FDA National Drug Classification (NDC) codes in the Medicare linked claims data.

After all exclusions, 2,543 total individuals remained in the analytic dataset. The process of exclusions is shown in **Fig S1**. Metformin and DPP4i use was classified by the presence of ≥2 NDC codes (**Table S2**) greater than or equal to 14 days apart. Individuals with no claims following date of diagnosis were excluded from the analysis. Demographics of the final dataset stratified by treatment group are shown in **Table S3**.

Data processing and analyses were conducted in SAS 9.4 and R 4.3 using the following packages: using the following packages: gtsummary, patchwork, survival, survminer, and tidyverse suite.

### Bioinformatic analysis

#### DPP-4 expression in single cell-RNA datasets

Raw data was downloaded from the Gene Expression Omnibus (GEO) under accession number GSE11789^102^ and processed as described before^103^. Briefly, the data were normalized for library size and log-transformed.

Data matrices were imputed using a Markov affinity-based graph approach^104^ to mitigate drop-out effects. Non-tumor cell types were annotated using the Mann-Whitney-Wilcoxon gene set test (mww-GST)^105^, based on a panel of 295 immune and stromal gene signatures^103^. Expression levels of DPP4 were assessed across immune cell populations. The analyses were performed using R (version 4.2.1).

#### Differential gene expression and pathway enrichment Analysis of DPP4-Associated Programs

Single-cell RNA sequencing data were obtained from GBMap, a comprehensive and harmonized atlas of GBM samples to identify which cells expressed DPP4. For differential expression analysis, we focused specifically on the dataset GSE163108^51^ due to its uniquely high number of patient samples and selection for T cells processed within the same experimental batch. The GSE163108 dataset comprises over 31 glioma patients and thus the largest number of T cells in one study. Prior to differential expression analysis, cells were classified as DPP4-positive or DPP4-negative based on whether normalized DPP4 expression exceeded zero. Only pseudo-bulk samples with total UMI counts above 5,000 were retained. Raw counts were aggregated into pseudo-bulk profiles using the AggregateExpression function from the *Seurat* R package, grouping cells by donor ID and DPP4 expression status. Differential gene expression analysis was performed using the R package *limma* . A design matrix was created incorporating DPP4 status and donor identity (∼ DPP4 _group + donor_id) to account for inter-individual variability. Count data were transformed using voom() to model the meanvariance trend. Linear modeling and empirical Bayes moderation were conducted using lmFit() and eBayes(). Gene set enrichment analysis was performed using the *fgsea* and *MsigDB* R packages, using logFC values to rank genes and KEGG ontologies as pathways. Enrichment was performed using 10,000 permutations. Pathways with an adjusted p-value < 0.05 were considered significantly enriched. The top 15 pathways, based on normalized enrichment score (NES), were visualized using bar plots.

## Data analysis

Flow cytometry data was acquired on LSR-Fortessa-HTS and Cytek Aurora or a Beckman Coulter CytoFlex LX flow cytometer. FACS data was analyzed using FlowJo software 10.8.

## Acknowledgements

This work is supported by V Scholar Award V2024-033 (D.B.), Sylvester Comprehensive Cancer Center (SCCC) Start-up Fund (D.B., D.C.W.), NCI R00 CA248611 (D.B.), NCI R01 CA253986 (D.B.L.), NCI R00 CA277242 (D.C.W.), the Danish Medical Research Council (B.W.K.), Lundbeck Foundation (B.W.K.), and NCI R37 CA285434 (Z.A.B.). Research reported in this publication was performed in part at the Flow Cytometry Shared Resource (FCSR; RRID: SCR_022501) of the SCCC at the University of Miami Miller School of Medicine, which is supported by the National Cancer Institute Cancer Center Support Grant (CCSG) P30-CA240139. SCCC also supported some of the metabolomic studies. This project was further supported with pilot funding from the Duke Center for Brain and Spine Metastasis (Q.T.O) and grants from the Preston Robert Tisch Brain Tumor Center, the Duke Cancer Institute (P30CA014236) (Q.T.O). The authors also thank the Human Immunology Core at the Perelman School of Medicine at the University of Pennsylvania for assistance with healthy donor PBMCs for CAR T cell generation. The HIC is supported in part by NIH P30 AI045008 and P30 CA016520. HIC RRID: SCR_022380. This study used the linked SEER-Medicare database. The interpretation and reporting of these data are the sole responsibility of the authors. The authors acknowledge the efforts of the National Cancer Institute; Information Management Services (IMS), Inc.; and the Surveillance, Epidemiology, and End Results (SEER) Program tumor registries in the creation of the SEER-Medicare database. The collection of cancer incidence data used in this study was supported by the California Department of Public Health pursuant to California Health and Safety Code Section 103885; Centers for Disease Control and Prevention’s (CDC) National Program of Cancer Registries, under cooperative agreement 1NU58DP007156; the National Cancer Institute’s Surveillance, Epidemiology and End Results Program under contract HHSN261201800032I awarded to the University of California, San Francisco, contract HHSN261201800015I awarded to the University of Southern California, and contract HHSN261201800009I awarded to the Public Health Institute. The ideas and opinions expressed herein are those of the author(s) and do not necessarily reflect the opinions of the State of California, Department of Public Health, the National Cancer Institute, and the Centers for Disease Control and Prevention or their Contractors and Subcontractors.

---

**Supplemental Figure 1.**
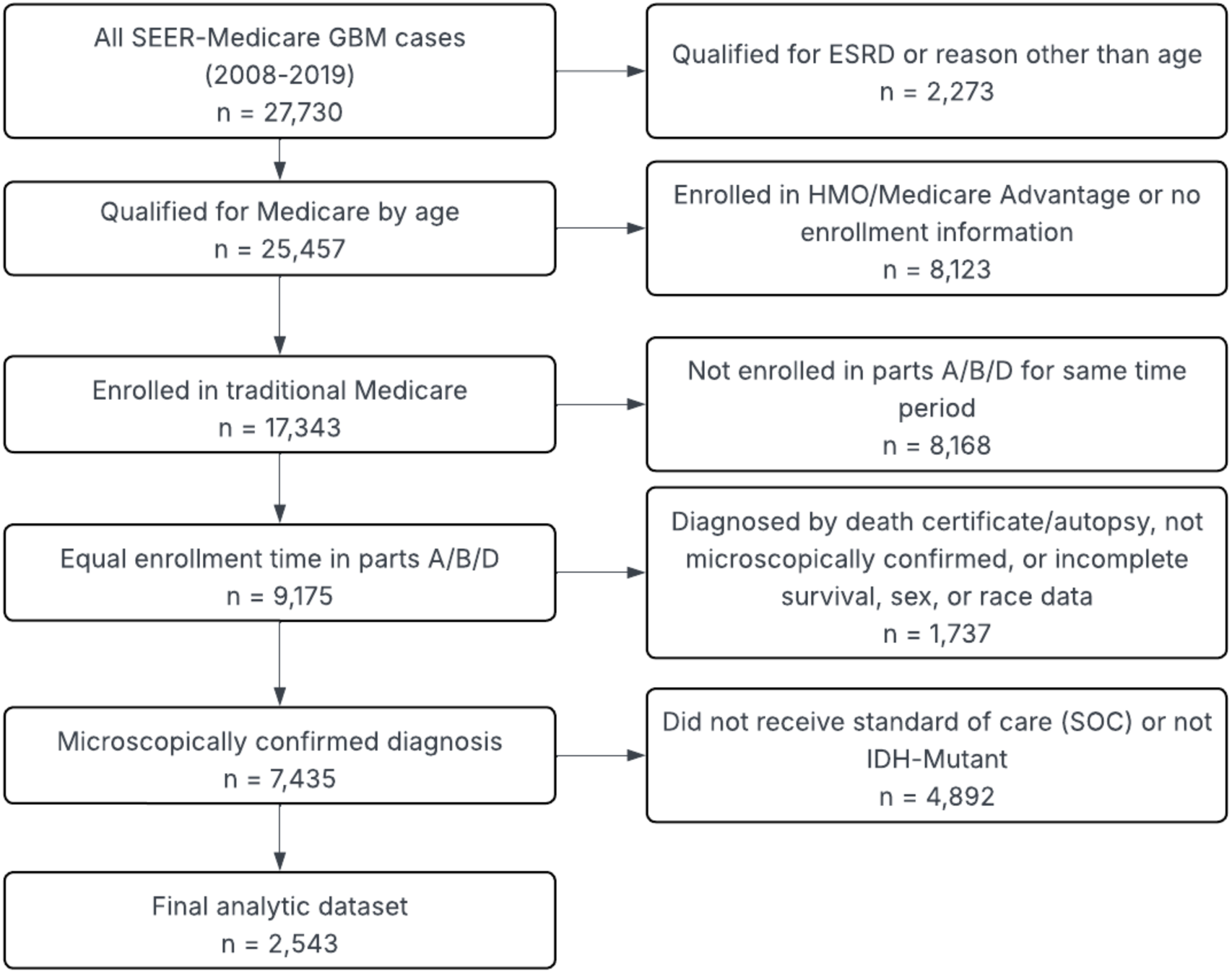
Diagram of SEER-Medicare GBM analysis. Workflow of the inclusion criteria to define GBM patients for subsequent survival analysis.

**Supplemental Figure 2.**
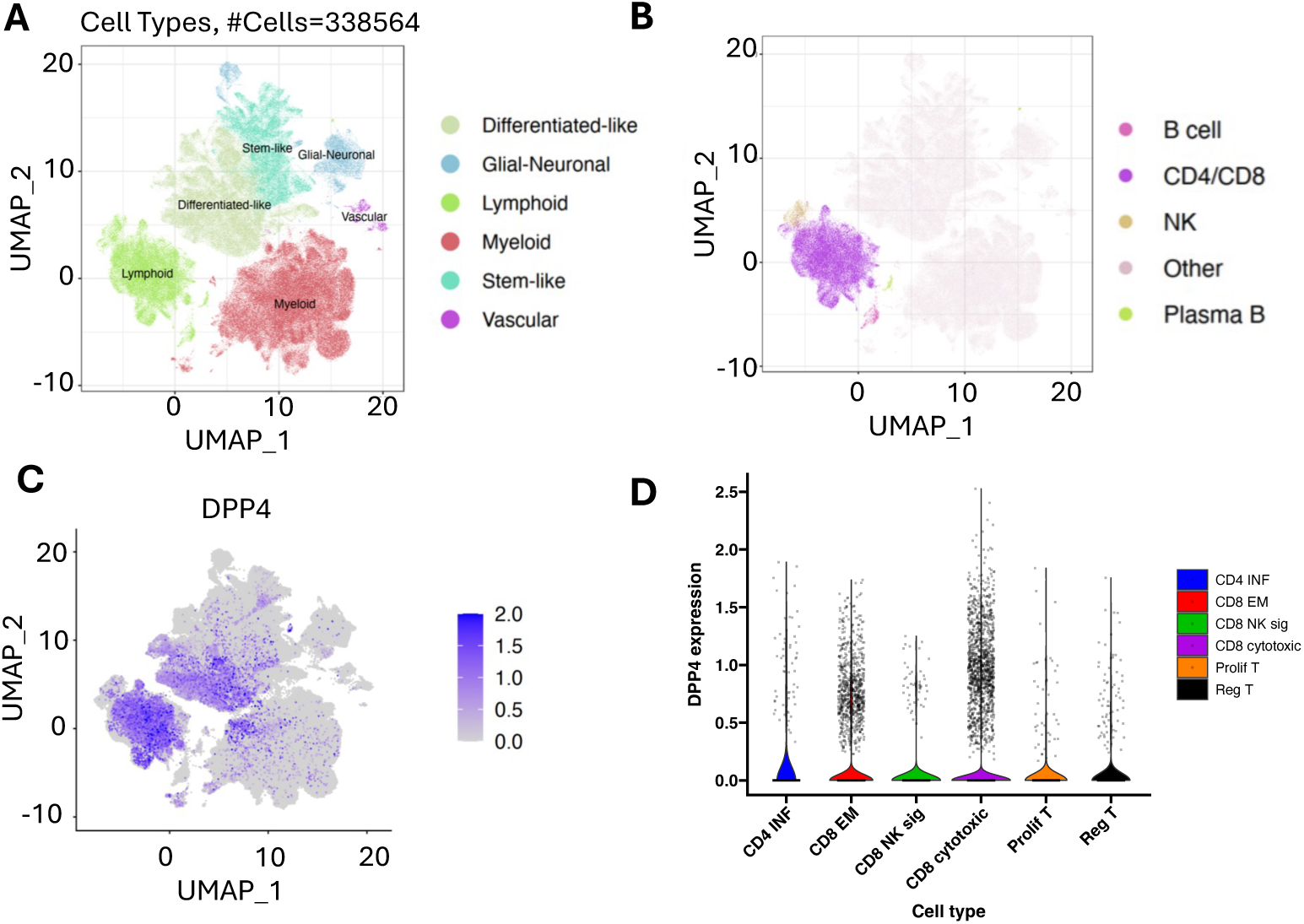
GBM-infiltrating T cells express high levels of DPP-4. GBMap core (> 300,000 cells total) was used to determine clusters in single-cell RNA-seq data set (GSE163108). UMAP graphs demonstrating **(A)** major cell types, **(B)** lymphocytes, and **(C)** *DPP4* expression within these populations. **(D)**. Expression of DPP4 across lymphocyte clusters.

**Supplemental Figure 3.**
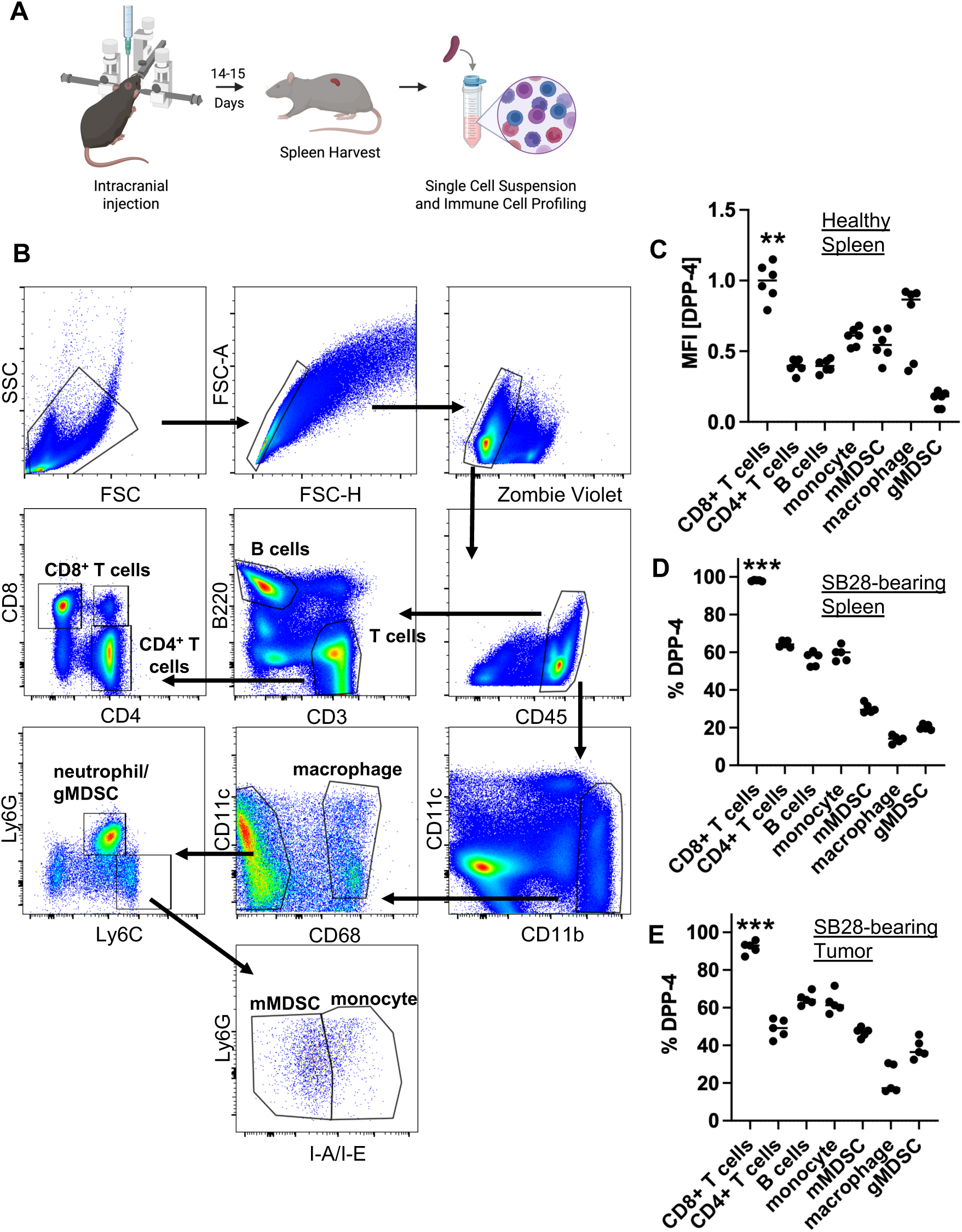
DPP-4 is differentially expressed by immune populations. **(A)** Diagram depicting intracranial injection protocol for immunophenotyping. **(B)** Gating strategy to identify immune populations from mouse spleens and tumors**. (C)** Mean fluorescence intensity (MFI) of DPP-4 from spleens of healthy mice. n=6; ** p<0.01 by two-way ANOVA. Percent DPP-4 expression in **(D)** splenic and **(E)** tumor-infiltrating immune populations. n=5; *** p<0.001 by one-way ANOVA.

**Supplemental Figure 4.**
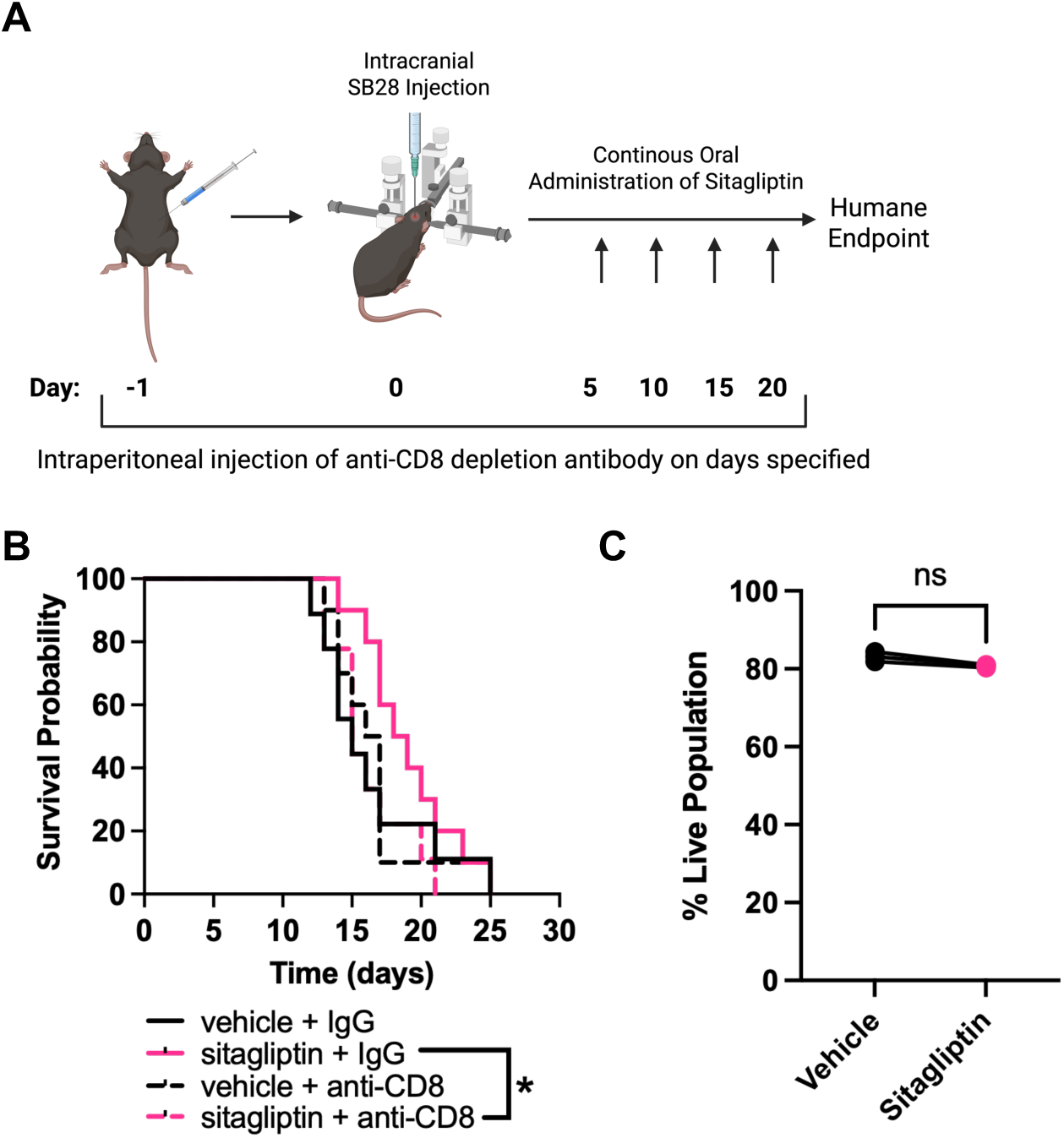
Sitagliptin acts on CD8+ T-Cells to improve GBM outcomes. **(A)** C57BL/6 male mice were orthotopically implanted with 20,000 SB28 and treated with 4 g/L sitagliptin in water bottle starting on Day 0. Mice were intraperitoneally injected with 200 μg anti-CD8 or IgG2b isotype control antibodies on Days -1 and 0 and then with 100 μg every 5 days post intracranial injection until humane endpoint. **(B)** Kaplan-Meier curve of SB28-bearing mice treated vehicle/sitagliptin + anti-CD8/IgG2b. n=10/group combined from 2-independent experiments. **(C)** SB28 were treated with sitagliptin at 100 μM for two days and live population frequency evaluated by flow cytometry. n=3; ns=not significant by t-test.

**Supplemental Figure 5.**
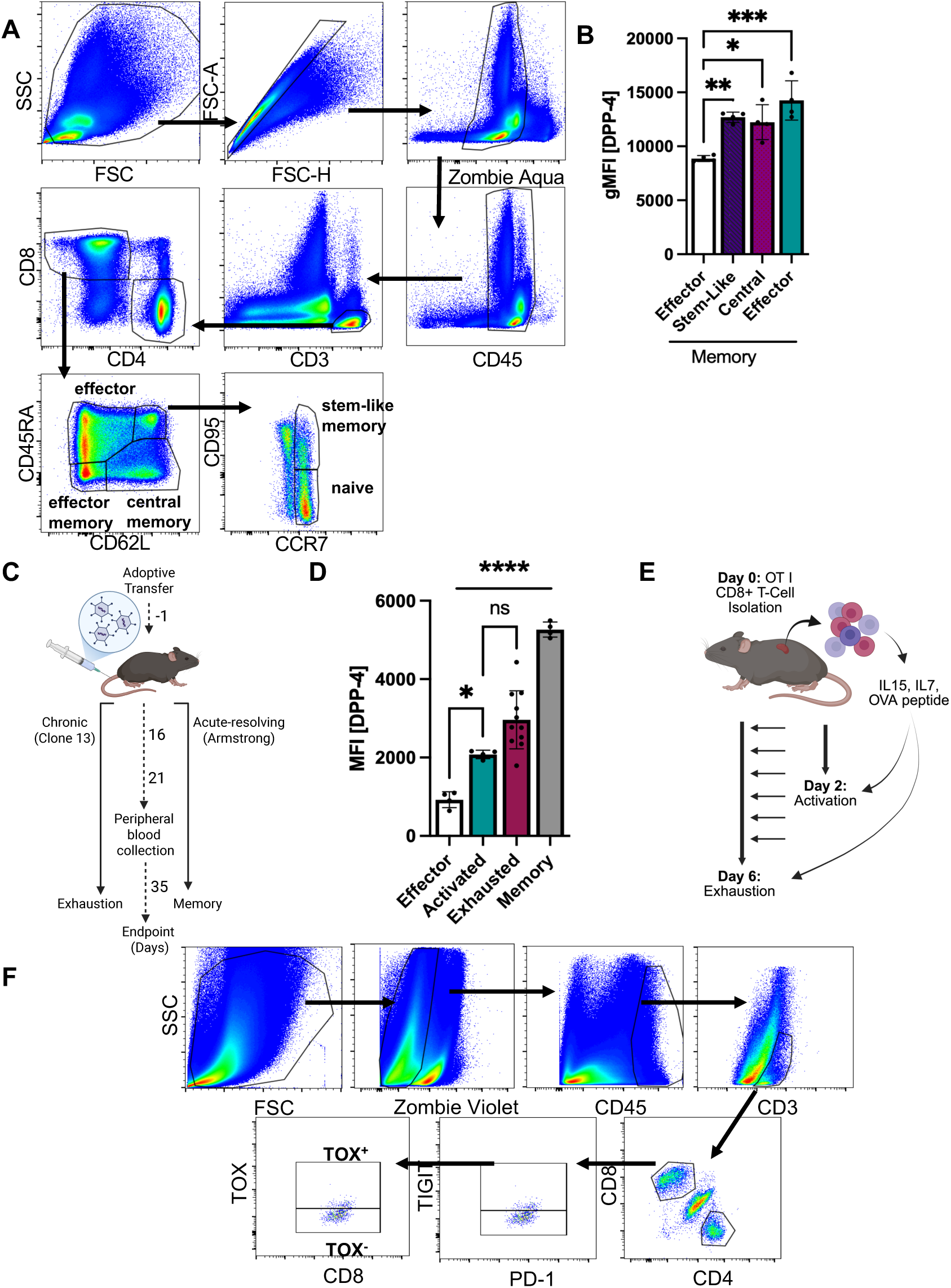
DPP-4 expression is informed by CD8^+^ T cell states. **(A)** Gating strategy to identify immune populations from healthy human PBMCs. **(B)** DPP-4 geometric MFI in memory CD8^+^ T cells from healthy donors. n=4; * p<0.05, ** p<0.01, *** p<0.001 by ordinary one-way ANOVA. **(C)** Diagram showing how naïve CD45.1^+^ p14 CD8^+^ T cells were adoptively transferred into recipient wild-type CD45.2^+^ mice. Recipient mice were infected with 2 x 10^5^ plaque forming units (PFU) of LCMV Armstrong intraperitoneally (i.p.) or with 4 x 10^6^ LCMV clone 13 intravenously (i.v.) the day after. Peripheral blood was collected on day 16 and 21 to confirm infection, and spleens collected on day 35 for endpoint immune analysis**. (D)** DPP-4 MFI from splenic CD8^+^ T cells infected with two different LCMV strains to promote effector, activated, exhausted, and memory differentiation. Clone 13, n=11, Amstrong, n=4; * p<0.05, **** p<0.0001, ns=not significant by one-way ANOVA. **(E)** Splenic CD8^+^ T cells from OT-I transgenic mice were activated with 10 ng OVA for two days or repetitively stimulated for five days to drive exhaustion. **(F)** Gating strategy used to define exhausted CD8^+^ T cells within human high-grade gliomas and surrounding tissue.

**Supplemental Figure 6.**
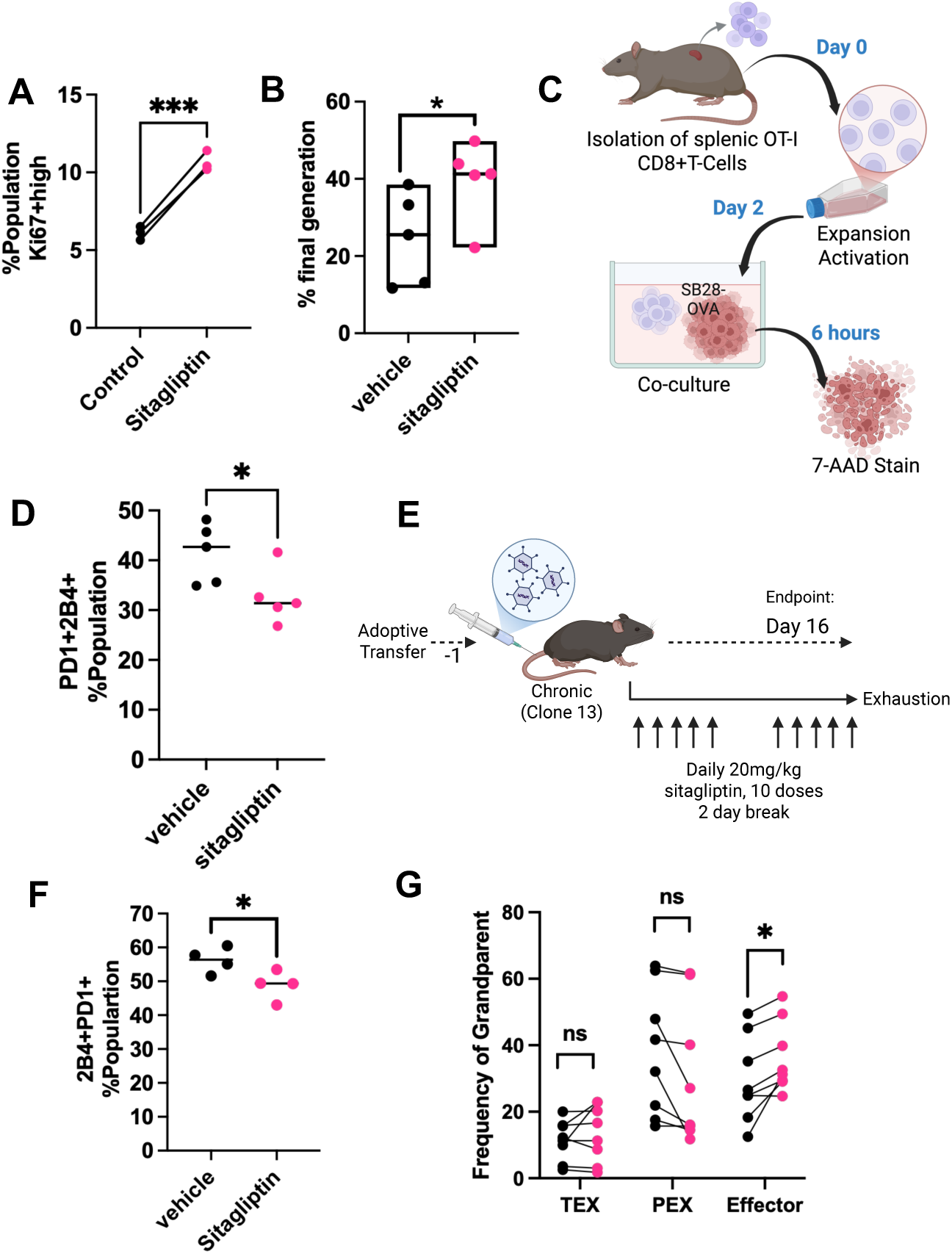
Sitagliptin treatment promotes CD8^+^ T cell effector function and delays exhaustion. **(A)** Mouse CD8^+^ T cells activated with anti-CD3/CD28 plus 100IU IL-2 in the presence of 100 μM sitagliptin or vehicle control for 4 days were intracellularly stained with Ki67 to determine proliferative capacity. n=4; *** p<0.001 by paired t-test. **(B)** Human CD8^+^ T cells isolated from healthy PBMCs were expanded with anti-CD3/CD28 and 100IU IL-2 for 6 days, and CFSE dilution determined by flow cytometry. n=5; * p<0.05 by paired t-test**. (C)** Schematics of the killing assay. Splenic CD8^+^ T cells from transgenic OT-I mice were isolated and activated with 10 ng OVA for 2 days, co-cultured at 1:1 E:T with either OVA- or empty vector-expressing cancer cells for six hours and then stained with apoptotic marker 7-AAD to determine cancer cell death by flow cytometry. **(D)** Mouse CD8^+^ T cells activated with anti-CD3/CD28 plus 100 IU IL-2 in the presence of 100 μM sitagliptin or vehicle control for 4 days were stained with exhaustion marker, anti-PD-1 and anti-2B4, to determine exhaustion post treatment. n=5; * p<0.05 by paired t-test. **(E)** Schematics of the LCMV experiment. Naïve CD45.1^+^ p14 CD8^+^ T cells were adoptively transferred into recipient wild-type CD45.2^+^ mice, and then recipient mice were infected with 4 x 10^6^ PFU LCMV clone 13 intravenously (i.v.) the day after. Recipient mice were treated intraperitoneally with 20 mg/kg sitagliptin daily for 5 continues doses, 2-day break, and 5 more doses for a total of 10 doses. Mice were then observed daily for indications of endpoint. **(F)** Splenic CD8^+^ T cells isolated from **(E)** at endpoint and stained with exhaustion marker, anti-PD-1 and anti-2B4, to determine sitagliptin induced effect in LCMV Clone 13 model. n=4; * p<0.05 by t-test. **(G)** Frequency of grandparents for TEX, PEX, and effector populations was determined for OT-I CD8^+^ T cells repetitively stimulated with 10 ng OVA for 5 days. n=7; * p<0.05, ns=not significant by paired t-test.

**Supplemental Figure 7.**
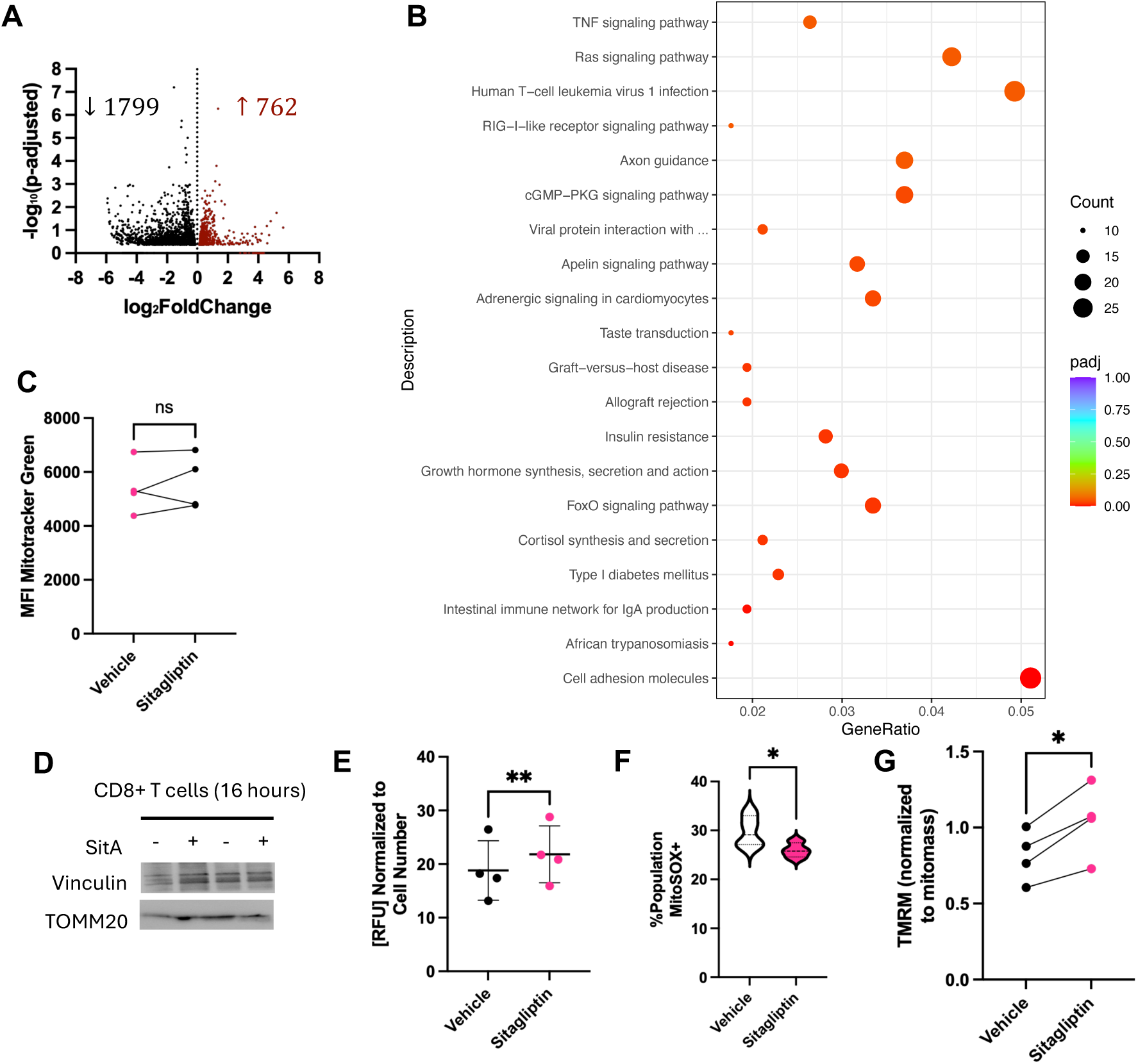
Sitagliptin reprograms CD8^+^ T cells. **(A)** Volcano plot of differentially regulated genes (1799 down and 762 up) upon sitagliptin treatment for 4 days. **(B)** Top differentially down-regulated pathways in sitagliptin-treated CD8^+^ T cells based on genes expressed >2-fold with p<0.05. n=6. **(C)** Splenic mouse CD8^+^ T cells were activated with anti-CD3/CD28 plus 100 IU/mL IL-2 in the presence of 100 μM sitagliptin or vehicle control for 2 days and then stained with 20 nM of the mitochondrial permeable stain, Mitotracker green, for 30 minutes to quantify mitochondria mass. n=4; ns=not significant by paired t-test. **(D)** Splenic mouse CD8^+^ T cells were activated with anti-CD3/CD28 plus 100 IU/mL IL-2 in the presence of 100 μM sitagliptin or vehicle control overnight. Cells were collected and lysed with RIPA buffer containing phosphatase/protease inhibitors, and 20 μM of protein were loaded on an SDS-Page for protein resolution, and western blotting analysis. **(E)** Splenic mouse CD8^+^ T cells were activated with anti-CD3/CD28 plus 100 IU/mL IL-2 in the presence of 100 μM sitagliptin or vehicle control for 2 days, and then ATP levels were determined with CellTiter-Glo Luminescent Cell Viability Assay. n=4; ** p<0.01 by paired t-test. **(F)** CD8^+^ T cells activated for two days were incubated with the superoxide indicator, MitoSOX Deep Red, at 500 nM for 60 minutes. The percentage of T cells positive for superoxide accumulation was determined by flow cytometry. n=4; * p<0.05 by paired t-test. **(G)** CD8^+^ T cells from OT-I mice were isolated and stimulated with 10 ng OVA repetitively for 5 days to elicit exhaustion. On Day 6, cells were incubated with Mitotracker and TMRM. TMRM MFI was normalized to mitochondrial mass. n=4; * p<0.05 by paired t-test.

**Supplemental Figure 8.**
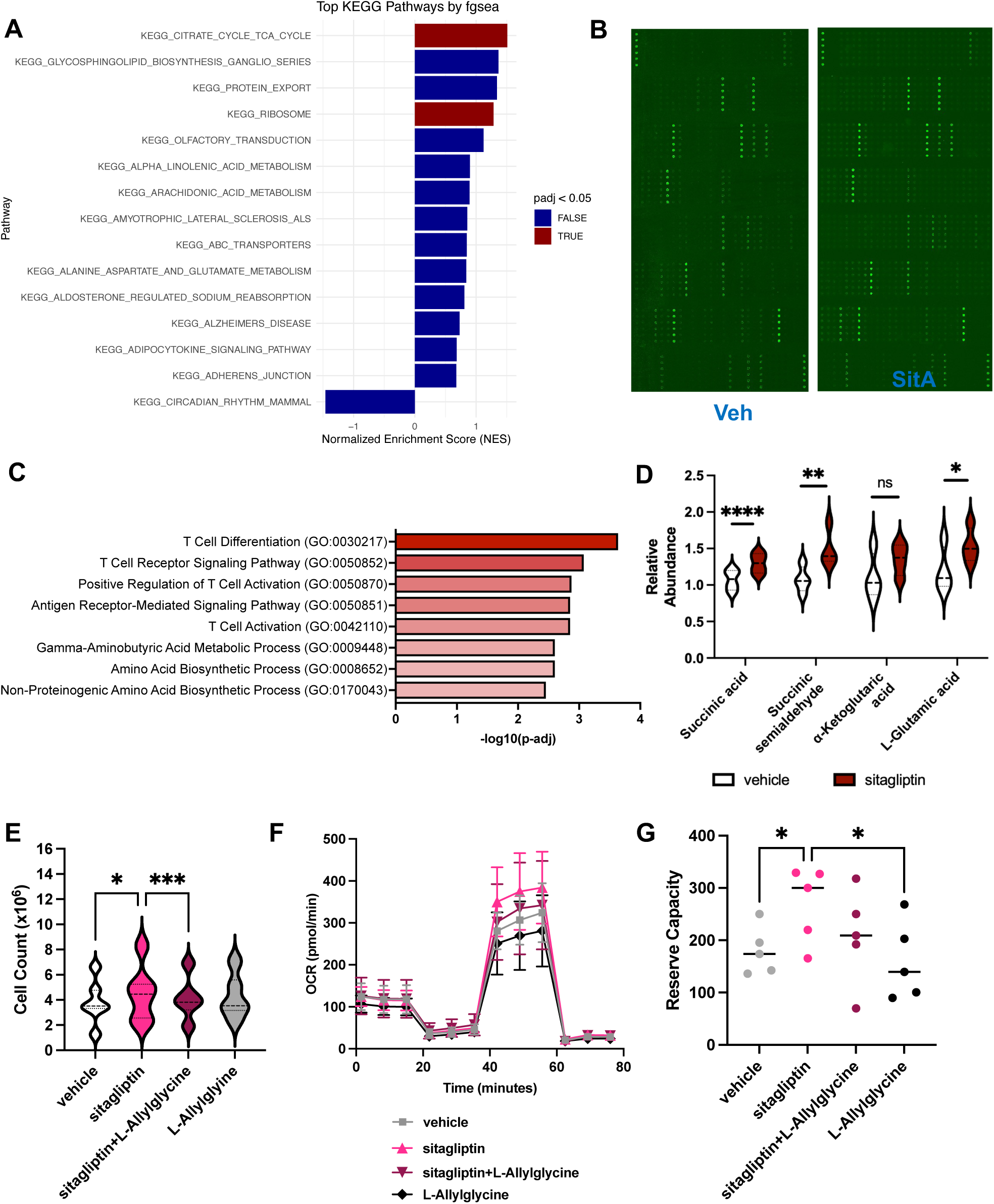
DPP-4 dictates metabolic changes in CD8^+^ T cells. **(A)** Top differentially altered pathways in DPP4^high^ vs. DPP4^low^ human GBM-infiltrating CD8^+^ T cells. Pathways with an adjusted p-value < 0.05 were considered significantly enriched. **(B)** Phospho-proteomic array fluorescent signal from splenic CD8^+^ T cells pooled from 4 biological replicates and activated overnight with anti-CD3/CD28 plus 100 IU/mL IL-2 with 100 μM sitagliptin or vehicle control. **(C)** Differentially regulated pathways from **(B)**. **(D)** Targeted metabolomics was performed with mouse CD8^+^ T cells activated for two days with 100 μM sitagliptin or vehicle control. Metabolite quantification for GABA shunt pathway intermediates was determined. n=4; * p<0.05, ** p<0.01, **** p<0.0001, ns=not significant by paired t-test. **(E)** Human CD8^+^ T cells from healthy human donors were isolated form PBMCs and activated with anti-CD3/CD28 plus 100 IU/mL IL-2 in the presence of sitagliptin (100 μM), the GAD1 inhibitor, L-Allylglycine (100 μM), or combination treatment for 6 days. Cells were then collected, stained with trypan blue, and quantified using an automatic cell counter. n=4; * p<0.05, *** p<0.001 by mixed effect analysis. Human CD8^+^ T cells from healthy human donors were isolated from PBMCs and expanded with anti-CD3/CD28 plus 100 IU/mL IL-2 for 6 days to reduce donor variability. Cells were then further activated in the presence of sitagliptin (100 μM), L-Allylglycine (200 μM), or combination treatment for four days. Seahorse T-Cell Fitness Assay was performed to determine **(F)** OCR and **(G)** reserve capacity. n=6; * p<0.05 by Friedman test.

**Supplemental Figure 9.**
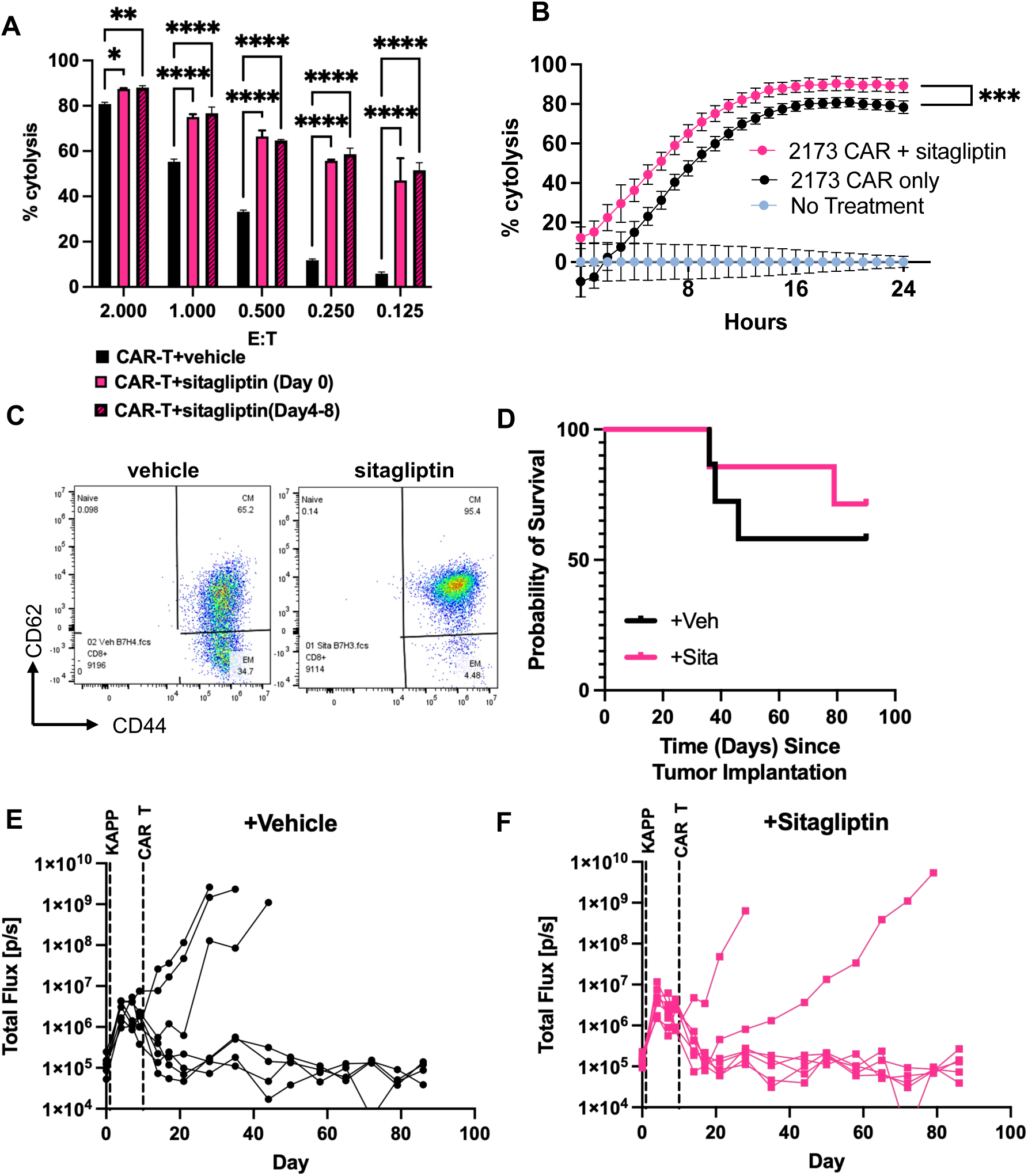
Sitagliptin treatment enhances CAR T-cell cytotoxic function. **(A)** Percent lysis of OCI-LY1, a human diffuse large B-cell lymphoma (DLBCL) cell line, co-cultured with CD19-targetting CAR-T manufactured with sitagliptin or vehicle control at different E:T Ratios **(B)** Percent killing of U87-EGFRvIII cells incubated with IL-13Rα2 CAR T-cells for 24 hours. **(C)** Dot plot of B7-H3 CAR T cell phenotyping based on CD44 and CD62 expression on day 5 after expansion with 50 U/mL of IL-2 and 100 μM of sitagliptin or vehicle control. **(D)** Kaplan-Meier curve of mice intracranially injected with the syngeneic mouse cell line, KAPP, and treated with CAR-T cells. Total flux plots of mice from **(D)** treated with one dose of either **(E)** vehicle or **(D)** sitagliptin-treated B7-H3-targetting CAR-T cells on day 10 post intracranial injection.

**Table S1.** Antibody information.

| Target | Fluorophore | Reactivity | Catalog | Clone | Manufacturer | Dilution |
| --- | --- | --- | --- | --- | --- | --- |
| CD127 | PE-Cy5 | Mouse | A7R34 | 135015 | Biolegend | 1:500 |
| CD160 | PE | Mouse | 7H1 | 143003 | Biolegend | 1:200 |
| CD19 | APC-Cy7 | Human | HIB19 | 302217 | Biolegend | 1:100 |
| CD197 (CCR7) | APC | Human | G043H7 | 353213 | Biolegend | 1:100 |
| CD223 (LAG3) | BV650 | Human | 11C3C65 | 369307 | Biolegend | 1:200 |
| CD244.2 | APC | Mouse | m2b4<br>(B6)458.1 | 133507 | Biolegend | 1:200 |
| CD26 (DPP-4) | PE-Cy7 | Human | BA5b | 302713 | Biolegend | 1:100 |
| CD26 (DPP-4) | FITC | Mouse | H194-112 | 137805 | Biolegend | 1:100 |
| CD3 | PerCP-Cy5.5 | Human | SK7 | 344807 | Biolegend | 1:100 |
| CD39 | BUV395 | Mouse | Y23-1185 | 567264 | BD Biosciences | 1:100 |
| CD39 | AF700 | Mouse | 24DMS1 | 56-0391-82 | Invitrogen | 1:100 |
| CD39 | PerCP-eF710 | Mouse | 24DMS1 | 46-0391-82 | Invitrogen | 1:50 |
| CD4 | BUV805 | Mouse | GK1.5 | 612900 | BD Biosciences | 1:500 |
| CD4 | APC-Cy7 | Mouse | GK1.5 | 552051 | BD Biosciences | 1:50 |
| CD4 | AF700 | Mouse | GK1.5 | 100429 | Biolegend | 1:200 |
| CD4 | BV605 | Human | SK3 | 344645 | Biolegend | 1:100 |
| CD44 | BUV395 | Mouse | IM7 | 568507 | BD Biosciences | 1:500 |
| CD44 | PE | Mouse | IM7 | 553134 | BD Biosciences | 1:50 |
| CD44 | BV510 | Mouse | IM7 | 103043 | Biolegend | 1:300 |
| CD44 | BV785 | Mouse | IM7 | 103041 | Biolegend | 1:300 |
| CD45 | AF700 | Human | HI30 | 304023 | Biolegend | 1:100 |
| CD45.1 | BUV615 | Mouse | A20 | 751467 | BD Biosciences | 1:200 |
| CD45.1 | AF700 | Mouse | A20 | 110723 | Biolegend | 1:100 |
| CD45.1 | APC-Cy7 | Mouse | A20 | 110715 | Biolegend | 1:200 |
| CD45.2 | BV785 | Mouse | 104 | 109839 | Biolegend | 1:200 |
| CD45RA | BV421 | Human | HI100 | 304129 | Biolegend | 1:100 |
| CD56 (NCAM) | PE-Dazzle594 | Human | 5.1H11 | 362543 | Biolegend | 1:100 |
| CD62L | PerCp-Cy5.5 | Mouse | MEL-14 | 560513 | BD Biosciences | 1:50 |
| CD62L | PE | Human | S21001C | 385103 | Biolegend | 1:100 |
| CD69 | BUV737 | Mouse | H1.2F3 | 612793 | BD Biosciences | 1:500 |
| CD69 | PE-Cy5 | Mouse | H1.2F3 | 104509 | Biolegend | 1:100 |
| CD69 | BV650 | Human | FN50 | 310933 | Biolegend | 1:100 |
| CD8 | FITC | Mouse | 53-6.7 | 553030 | BD Biosciences | 1:50 |
| CD8-FITC | FITC | Human | SK1 | 344703 | Biolegend | 1:100 |
| CD8a | BV650 | Mouse | 53-6.7 | 100741 | Biolegend | 1:400 |
| CD8a | BUV805 | Mouse | 53-6.7 | 368-0081-82 | Invitrogen | 1:500 |
| CD95 (fas) | BV785 | Human | DX2 | 305645 | Biolegend | 1:100 |
| CX3CR1 | APC/Fire 750 | Mouse | SA011F11 | 149039 | Biolegend | 1:100 |
| CX3CR1 | BV711 | Mouse | SA011F11 | 149031 | Biolegend | 1:500 |
| EOMES | eFluor450 | Mouse | Dan11mag | 48-4875-82 | Invitrogen | 1:200 |
| Ki-67 | BV605 | Mouse | 16A8 | 652413 | Biolegend | 1:100 |
| KLRG1 | BV605 | Mouse<br>Human | 2F1/KLRG1 | 138419 | Biolegend | 1:200 |
| KLRG1 | FITC | Mouse<br>Human | 2F1/KLRG1 | 138409 | Biolegend | 1:200 |
| Ly108 | Pacific Blue | Mouse | 330-AJ | 134608 | Biolegend | 1:100 |
| PD-1 | BV421 | Mouse | 29F.1A12 | 135217 | Biolegend | 1:100 |
| PD-1 | PE-Cy7 | Mouse | 29F.1A12 | 135215 | Biolegend | 1:200 |
| PD-1 | PE | Mouse | 29F.1A12 | 135205 |  |  |
| PD-1 | PE-Dazzle594 | Mouse | 29F.1A12 | 135227 | Biolegend | 1:400 |
| PD-1 | BV605 | Mouse | 29F.1A12 | 135220 | Biolegend | 1:50 |
| TCF-7/TCF-1 | PE | Mouse | S33-966 | 564217 | BD Biosciences | 1:200 |
| TIGIT | APC | Mouse | 1G9 | 142106 | Biolegend | 1:50 |
| TIGIT | BV421 | Mouse | 1G9 | 142111 | Biolegend | 1:50 |
| Tim-3 | BV421 | Mouse | B8.2C12 | 134019 | Biolegend | 1:50 |
| Tim-3 | BV605 | Mouse | RMT3-23 | 119721 | Biolegend | 1:100 |
| TOX | APC | Mouse<br>Human | REA473 | 130-129-208 | Miltenyi Biotec | 1:100 |

**Table S2.**
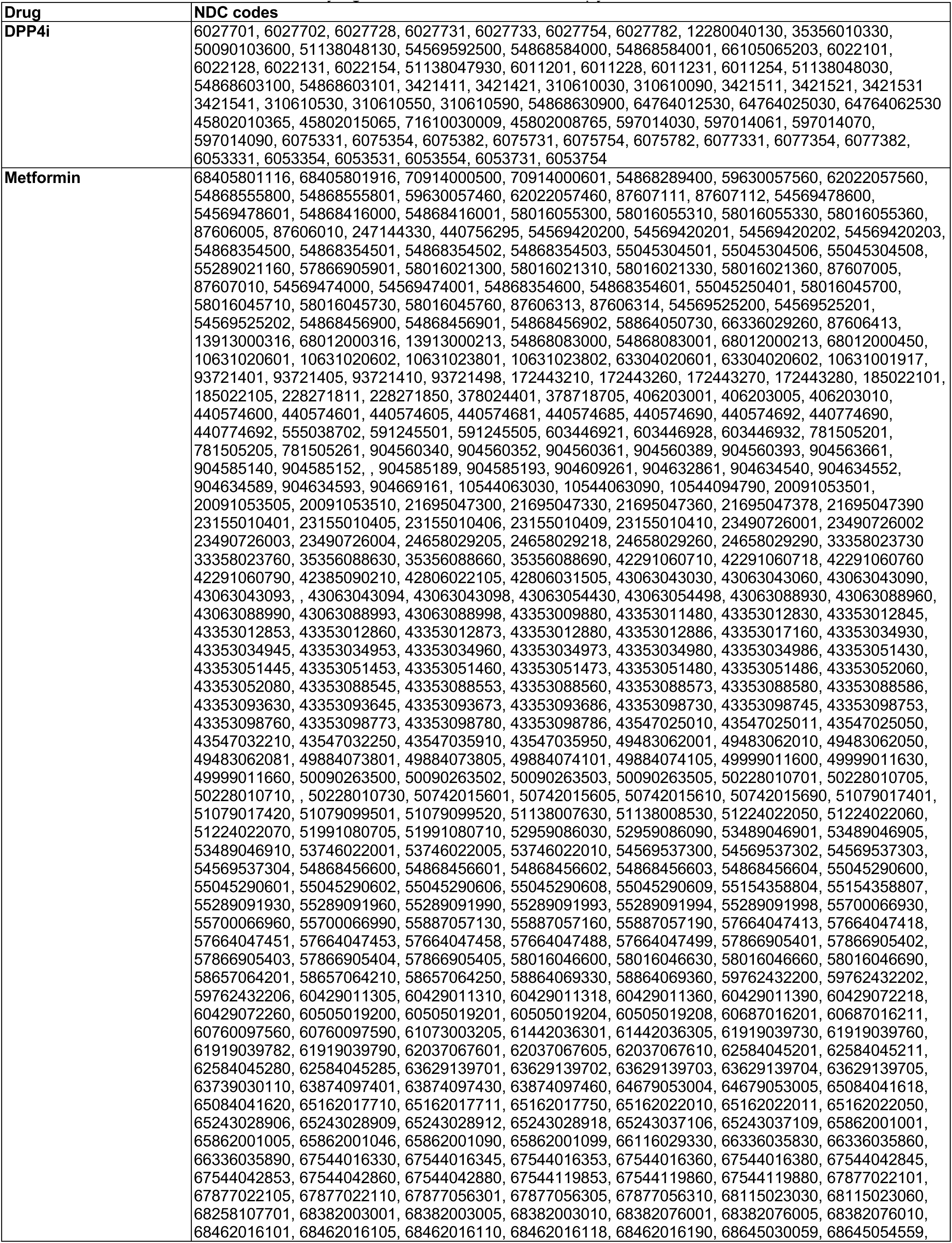
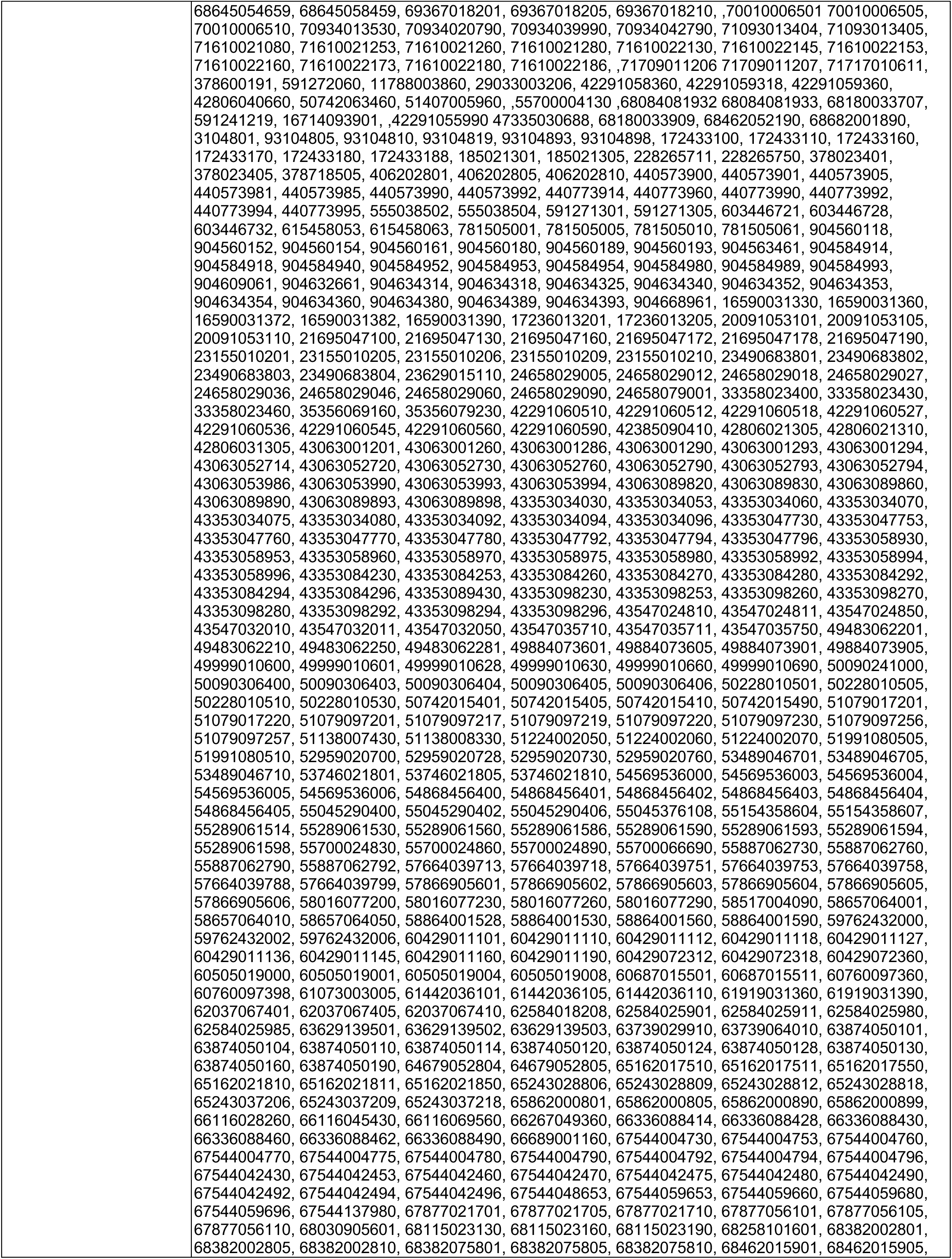
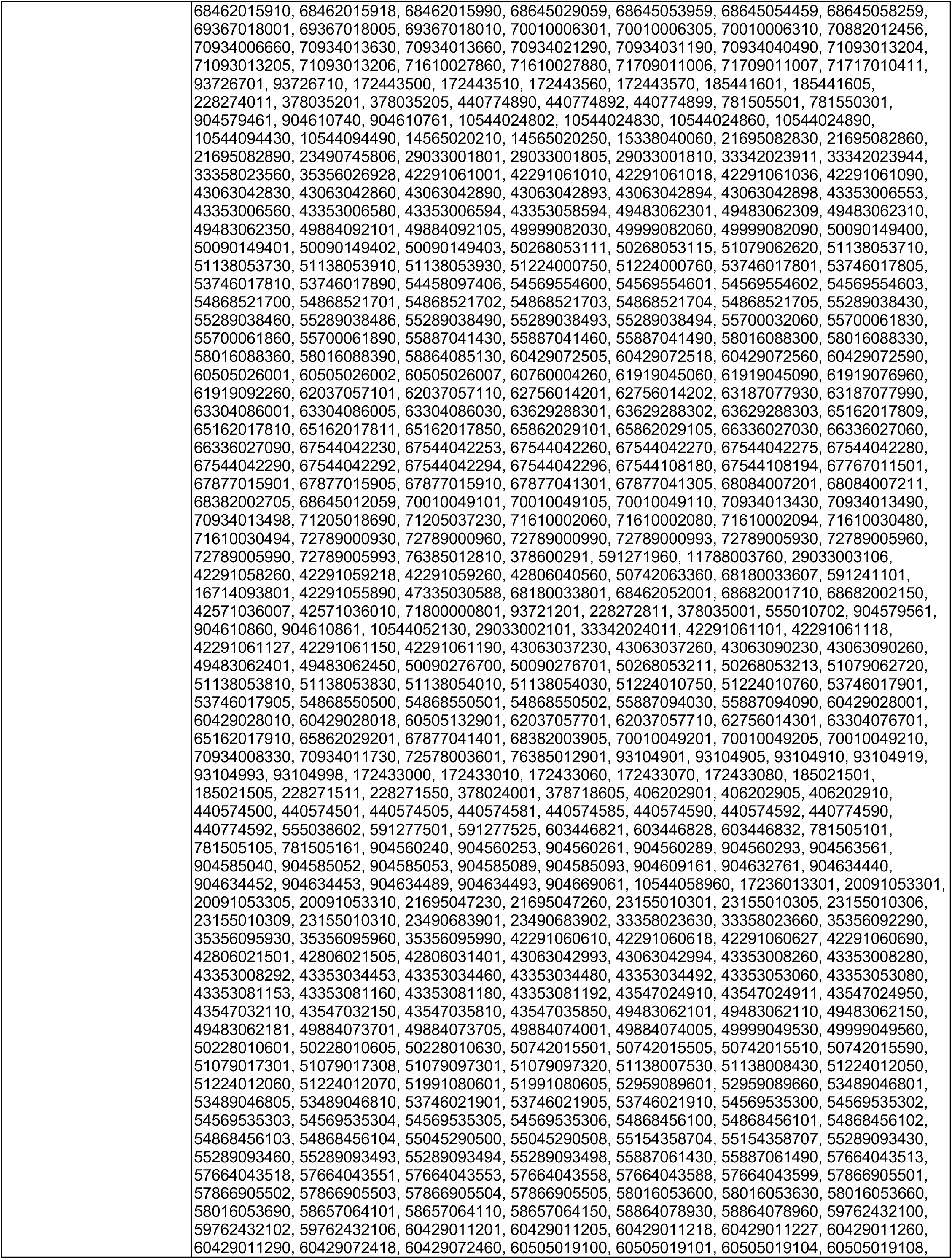
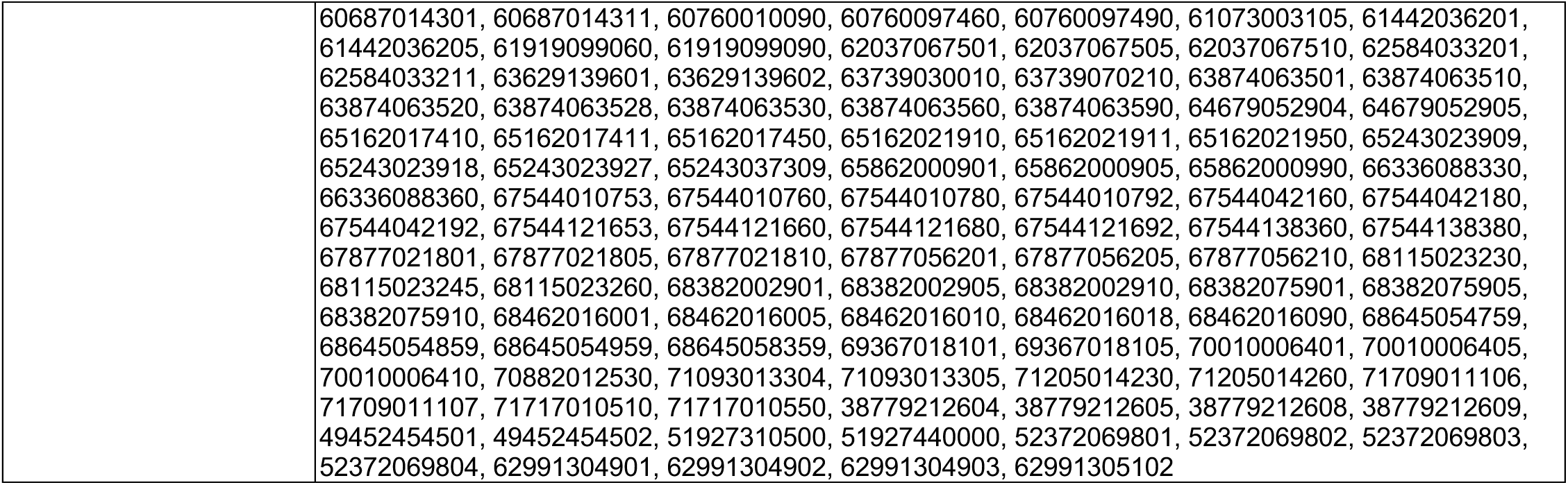
FDA NDC Codes for identifying DPP4i or metformin therapy use.

**Table S3.**
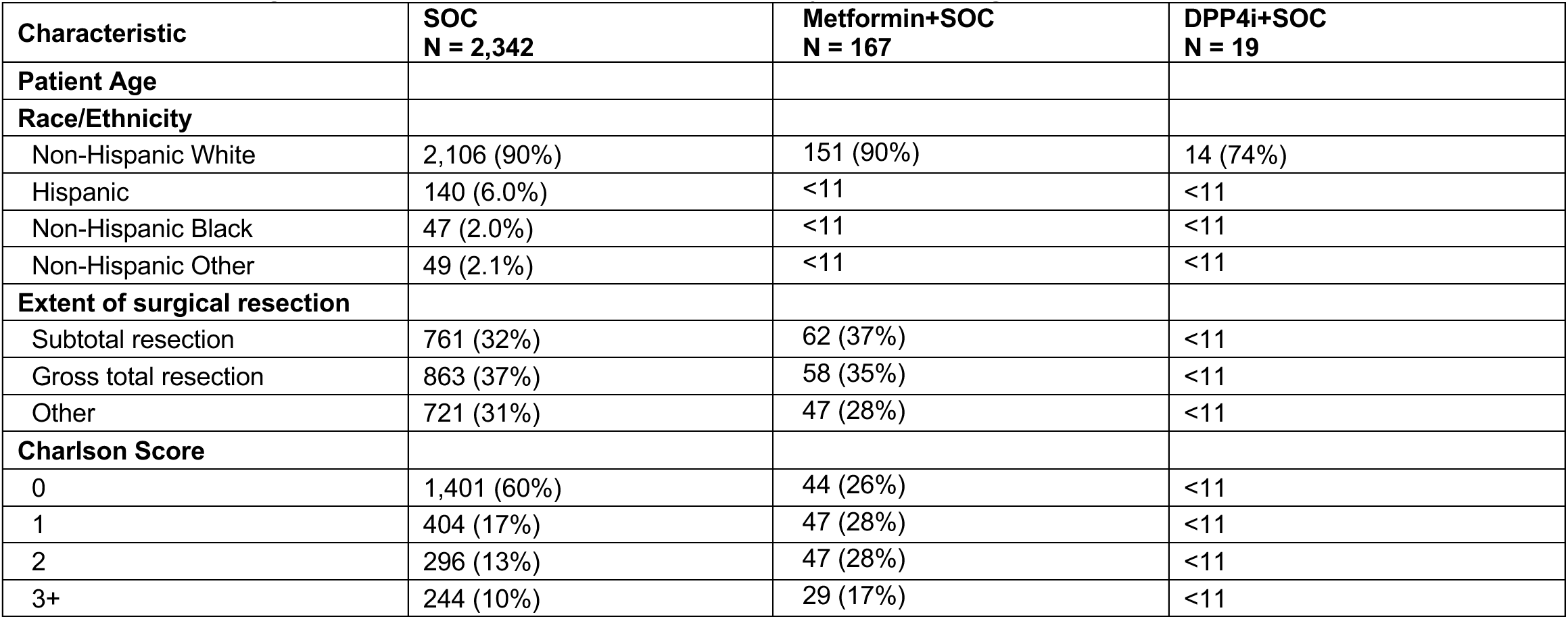
Demographics of included GBM patients by treatment group.

## References

1. Grywalska, E., Pasiarski, M., Gozdz, S., and Rolinski, J. (2018). Immune-checkpoint inhibitors for combating T-cell dysfunction in cancer. Onco Targets Ther 11, 6505–6524. 10.2147/OTT.S150817.

2. Belizario, J., and Destro Rodrigues, M.F. (2020). Checkpoint inhibitor blockade and epigenetic reprogrammability in CD8(+) T-cell activation and exhaustion. Ther Adv Vaccines Immunother 8, 2515135520904238. 10.1177/2515135520904238.

3. Blank, C.U., Haining, W.N., Held, W., Hogan, P.G., Kallies, A., Lugli, E., Lynn, R.C., Philip, M., Rao, A., Restifo, N.P., et al. (2019). Defining ’T cell exhaustion’. Nat Rev Immunol 19, 665–674. 10.1038/s41577-019-0221-9.

4. Philip, M., and Schietinger, A. (2022). CD8(+) T cell differentiation and dysfunction in cancer. Nat Rev Immunol 22, 209–223. 10.1038/s41577-021-00574-3.

5. Zhu, Y., Tan, H., Wang, J., Zhuang, H., Zhao, H., and Lu, X. (2024). Molecular insight into T cell exhaustion in hepatocellular carcinoma. Pharmacol Res 203, 107161. 10.1016/j.phrs.2024.107161.

6. Yu, Y.R., Imrichova, H., Wang, H., Chao, T., Xiao, Z., Gao, M., Rincon-Restrepo, M., Franco, F., Genolet, R., Cheng, W.C., et al. (2020). Disturbed mitochondrial dynamics in CD8(+) TILs reinforce T cell exhaustion. Nat Immunol 21, 1540–1551. 10.1038/s41590-020-0793-3.

7. Bengsch, B., Johnson, A.L., Kurachi, M., Odorizzi, P.M., Pauken, K.E., Attanasio, J., Stelekati, E., McLane, L.M., Paley, M.A., Delgoffe, G.M., and Wherry, E.J. (2016). Bioenergetic Insufficiencies Due to Metabolic Alterations Regulated by the Inhibitory Receptor PD-1 Are an Early Driver of CD8(+) T Cell Exhaustion. Immunity 45, 358–373. 10.1016/j.immuni.2016.07.008.

8. Scharping, N.E., Menk, A.V., Moreci, R.S., Whetstone, R.D., Dadey, R.E., Watkins, S.C., Ferris, R.L., and Delgoffe, G.M. (2016). The Tumor Microenvironment Represses T Cell Mitochondrial Biogenesis to Drive Intratumoral T Cell Metabolic Insufficiency and Dysfunction. Immunity 45, 374–388. 10.1016/j.immuni.2016.07.009.

9. Scharping, N.E., Rivadeneira, D.B., Menk, A.V., Vignali, P.D.A., Ford, B.R., Rittenhouse, N.L., Peralta, R., Wang, Y., Wang, Y., DePeaux, K., et al. (2021). Mitochondrial stress induced by continuous stimulation under hypoxia rapidly drives T cell exhaustion. Nat Immunol 22, 205–215. 10.1038/s41590-020-00834-9.

10. Simula, L., Fumagalli, M., Vimeux, L., Rajnpreht, I., Icard, P., Birsen, G., An, D., Pendino, F., Rouault, A., Bercovici, N., et al. (2024). Mitochondrial metabolism sustains CD8(+) T cell migration for an efficient infiltration into solid tumors. Nat Commun 15, 2203. 10.1038/s41467-024-46377-7.

11. Vardhana, S.A., Hwee, M.A., Berisa, M., Wells, D.K., Yost, K.E., King, B., Smith, M., Herrera, P.S., Chang, H.Y., Satpathy, A.T., et al. (2020). Impaired mitochondrial oxidative phosphorylation limits the self-renewal of T cells exposed to persistent antigen. Nat Immunol 21, 1022–1033. 10.1038/s41590-020-0725-2.

12. Wu, H., Zhao, X., Hochrein, S.M., Eckstein, M., Gubert, G.F., Knopper, K., Mansilla, A.M., Oner, A., Doucet-Ladeveze, R., Schmitz, W., et al. (2023). Mitochondrial dysfunction promotes the transition of precursor to terminally exhausted T cells through HIF-1alpha-mediated glycolytic reprogramming. Nat Commun 14, 6858. 10.1038/s41467-023-42634-3.

13. Guo, Y., Xie, Y.Q., Gao, M., Zhao, Y., Franco, F., Wenes, M., Siddiqui, I., Bevilacqua, A., Wang, H., Yang, H., et al. (2021). Metabolic reprogramming of terminally exhausted CD8(+) T cells by IL-10 enhances anti-tumor immunity. Nat Immunol 22, 746–756. 10.1038/s41590-021-00940-2.

14. Li, H., Zhao, A., Li, M., Shi, L., Han, Q., and Hou, Z. (2022). Targeting T-cell metabolism to boost immune checkpoint inhibitor therapy. Front Immunol 13, 1046755. 10.3389/fimmu.2022.1046755.

15. Zhang, H., Gao, J., Zhang, Z., and Zhang, B. (2025). Current status of chimeric antigen receptor T cell therapy and its exhaustion mechanism. Immunotherapy 17, 1039–1057. 10.1080/1750743X.2025.2560798.

16. Genoud, V., Marinari, E., Nikolaev, S.I., Castle, J.C., Bukur, V., Dietrich, P.Y., Okada, H., and Walker, P.R. (2018). Responsiveness to anti-PD-1 and anti-CTLA-4 immune checkpoint blockade in SB28 and GL261 mouse glioma models. Oncoimmunology 7, e1501137. 10.1080/2162402X.2018.1501137.

17. Omuro, A., Reardon, D.A., Sampson, J.H., Baehring, J., Sahebjam, S., Cloughesy, T.F., Chalamandaris, A.G., Potter, V., Butowski, N., and Lim, M. (2022). Nivolumab plus radiotherapy with or without temozolomide in newly diagnosed glioblastoma: Results from exploratory phase I cohorts of CheckMate 143. Neurooncol Adv 4, vdac025. 10.1093/noajnl/vdac025.

18. Lim, M., Weller, M., Idbaih, A., Steinbach, J., Finocchiaro, G., Raval, R.R., Ansstas, G., Baehring, J., Taylor, J.W., Honnorat, J., et al. (2022). Phase III trial of chemoradiotherapy with temozolomide plus nivolumab or placebo for newly diagnosed glioblastoma with methylated MGMT promoter. Neuro Oncol 24, 1935–1949. 10.1093/neuonc/noac116.

19. Koh, C.H., Lee, S., Kwak, M., Kim, B.S., and Chung, Y. (2023). CD8 T-cell subsets: heterogeneity, functions, and therapeutic potential. Exp Mol Med 55, 2287–2299. 10.1038/s12276-023-01105-x.

20. Prokhnevska, N., Cardenas, M.A., Valanparambil, R.M., Sobierajska, E., Barwick, B.G., Jansen, C., Reyes Moon, A., Gregorova, P., delBalzo, L., Greenwald, R., et al. (2023). CD8(+) T cell activation in cancer comprises an initial activation phase in lymph nodes followed by effector differentiation within the tumor. Immunity 56, 107–124 e105. 10.1016/j.immuni.2022.12.002.

21. Thompson, E.D., Enriquez, H.L., Fu, Y.X., and Engelhard, V.H. (2010). Tumor masses support naive T cell infiltration, activation, and differentiation into effectors. J Exp Med 207, 1791–1804. 10.1084/jem.20092454.

22. Steinert, E.M., Furtado Bruza, B., Danchine, V.D., Grant, R.A., Vasan, K., Kharel, A., Zhang, Y., Cui, W., Szibor, M., Weinberg, S.E., and Chandel, N.S. (2025). Mitochondrial respiration is necessary for CD8(+) T cell proliferation and cell fate. Nat Immunol 26, 1267–1274. 10.1038/s41590-025-02202-x.

23. Menk, A.V., Scharping, N.E., Moreci, R.S., Zeng, X., Guy, C., Salvatore, S., Bae, H., Xie, J., Young, H.A., Wendell, S.G., and Delgoffe, G.M. (2018). Early TCR Signaling Induces Rapid Aerobic Glycolysis Enabling Distinct Acute T Cell Effector Functions. Cell Rep 22, 1509–1521. 10.1016/j.celrep.2018.01.040.

24. Rangel Rivera, G.O., Knochelmann, H.M., Dwyer, C.J., Smith, A.S., Wyatt, M.M., Rivera-Reyes, A.M., Thaxton, J.E., and Paulos, C.M. (2021). Fundamentals of T Cell Metabolism and Strategies to Enhance Cancer Immunotherapy. Front Immunol 12, 645242. 10.3389/fimmu.2021.645242.

25. Ma, S., Ming, Y., Wu, J., and Cui, G. (2024). Cellular metabolism regulates the differentiation and function of T-cell subsets. Cell Mol Immunol 21, 419–435. 10.1038/s41423-024-01148-8.

26. Zheng, K., Zheng, X., and Yang, W. (2022). The Role of Metabolic Dysfunction in T-Cell Exhaustion During Chronic Viral Infection. Front Immunol 13, 843242. 10.3389/fimmu.2022.843242.

27. Gupta, S.S., Wang, J., and Chen, M. (2020). Metabolic Reprogramming in CD8(+) T Cells During Acute Viral Infections. Front Immunol 11, 1013. 10.3389/fimmu.2020.01013.

28. Liang, P., Li, Z., Chen, Z., Chen, Z., Jin, T., He, F., Chen, X., Gou, H., and Yang, K. (2025). Metabolic Reprogramming of Glycolysis, Lipids, and Amino Acids in Tumors: Impact on CD8+ T Cell Function and Targeted Therapeutic Strategies. FASEB J 39, e70520. 10.1096/fj.202403019R.

29. Wang, W., and Zou, W. (2020). Amino Acids and Their Transporters in T Cell Immunity and Cancer Therapy. Mol Cell 80, 384–395. 10.1016/j.molcel.2020.09.006.

30. Soto-Heredero, G., Desdin-Mico, G., and Mittelbrunn, M. (2021). Mitochondrial dysfunction defines T cell exhaustion. Cell Metab 33, 470–472. 10.1016/j.cmet.2021.02.010.

31. Cao, J., Liao, S., Zeng, F., Liao, Q., Luo, G., and Zhou, Y. (2023). Effects of altered glycolysis levels on CD8(+) T cell activation and function. Cell Death Dis 14, 407. 10.1038/s41419-023-05937-3.

32. Richter, F.C., Saliutina, M., Hegazy, A.N., and Bergthaler, A. (2024). Take my breath away-mitochondrial dysfunction drives CD8(+) T cell exhaustion. Genes Immun 25, 4–6. 10.1038/s41435-023-00233-8.

33. Lontos, K., Wang, Y., Joshi, S.K., Frisch, A.T., Watson, M.J., Kumar, A., Menk, A.V., Wang, Y., Cumberland, R., Lohmueller, J., et al. (2023). Metabolic reprogramming via an engineered PGC-1alpha improves human chimeric antigen receptor T-cell therapy against solid tumors. J Immunother Cancer 11. 10.1136/jitc-2022-006522.

34. Beckermann, K.E., Hongo, R., Ye, X., Young, K., Carbonell, K., Healey, D.C.C., Siska, P.J., Barone, S., Roe, C.E., Smith, C.C., et al. (2020). CD28 costimulation drives tumor-infiltrating T cell glycolysis to promote inflammation. JCI Insight 5. 10.1172/jci.insight.138729.

35. Hermans, D., Gautam, S., Garcia-Canaveras, J.C., Gromer, D., Mitra, S., Spolski, R., Li, P., Christensen, S., Nguyen, R., Lin, J.X., et al. (2020). Lactate dehydrogenase inhibition synergizes with IL-21 to promote CD8(+) T cell stemness and antitumor immunity. Proc Natl Acad Sci U S A 117, 6047–6055. 10.1073/pnas.1920413117.

36. Verma, S., Budhu, S., Serganova, I., Dong, L., Mangarin, L.M., Khan, J.F., Bah, M.A., Assouvie, A., Marouf, Y., Schulze, I., et al. (2024). Pharmacologic LDH inhibition redirects intratumoral glucose uptake and improves antitumor immunity in solid tumor models. J Clin Invest 134. 10.1172/JCI177606.

37. Wang, R., Dillon, C.P., Shi, L.Z., Milasta, S., Carter, R., Finkelstein, D., McCormick, L.L., Fitzgerald, P., Chi, H., Munger, J., and Green, D.R. (2011). The transcription factor Myc controls metabolic reprogramming upon T lymphocyte activation. Immunity 35, 871–882. 10.1016/j.immuni.2011.09.021.

38. Geiger, R., Rieckmann, J.C., Wolf, T., Basso, C., Feng, Y., Fuhrer, T., Kogadeeva, M., Picotti, P., Meissner, F., Mann, M., et al. (2016). L-Arginine Modulates T Cell Metabolism and Enhances Survival and Anti-tumor Activity. Cell 167, 829–842 e813. 10.1016/j.cell.2016.09.031.

39. Song, M., Sandoval, T.A., Chae, C.S., Chopra, S., Tan, C., Rutkowski, M.R., Raundhal, M., Chaurio, R.A., Payne, K.K., Konrad, C., et al. (2018). IRE1alpha-XBP1 controls T cell function in ovarian cancer by regulating mitochondrial activity. Nature 562, 423–428. 10.1038/s41586-018-0597-x.

40. Zhang, Y., Kurupati, R., Liu, L., Zhou, X.Y., Zhang, G., Hudaihed, A., Filisio, F., Giles-Davis, W., Xu, X., Karakousis, G.C., et al. (2017). Enhancing CD8(+) T Cell Fatty Acid Catabolism within a Metabolically Challenging Tumor Microenvironment Increases the Efficacy of Melanoma Immunotherapy. Cancer Cell 32, 377–391 e379. 10.1016/j.ccell.2017.08.004.

41. Bayik, D., Bartels, C.F., Lovrenert, K., Watson, D.C., Zhang, D., Kay, K., Lee, J., Lauko, A., Johnson, S., Lo, A., et al. (2022). Distinct Cell Adhesion Signature Defines Glioblastoma Myeloid-Derived Suppressor Cell Subsets. Cancer Res 82, 4274–4287. 10.1158/0008-5472.CAN-21-3840.

42. Deacon, C.F. (2019). Physiology and Pharmacology of DPP-4 in Glucose Homeostasis and the Treatment of Type 2 Diabetes. Front Endocrinol (Lausanne) 10, 80. 10.3389/fendo.2019.00080.

43. Omar, B., and Ahren, B. (2014). Pleiotropic mechanisms for the glucose-lowering action of DPP-4 inhibitors. Diabetes 63, 2196–2202. 10.2337/db14-0052.

44. Deacon, C.F. (2020). Dipeptidyl peptidase 4 inhibitors in the treatment of type 2 diabetes mellitus. Nat Rev Endocrinol 16, 642–653. 10.1038/s41574-020-0399-8.

45. Huang, J., Liu, X., Wei, Y., Li, X., Gao, S., Dong, L., Rao, X., and Zhong, J. (2022). Emerging Role of Dipeptidyl Peptidase-4 in Autoimmune Disease. Front Immunol 13, 830863. 10.3389/fimmu.2022.830863.

46. Wang, X., Zheng, P., Huang, G., Yang, L., and Zhou, Z. (2018). Dipeptidyl peptidase-4(DPP-4) inhibitors: promising new agents for autoimmune diabetes. Clin Exp Med 18, 473–480. 10.1007/s10238-018-0519-0.

47. Elmansi, A.M., Awad, M.E., Eisa, N.H., Kondrikov, D., Hussein, K.A., Aguilar-Perez, A., Herberg, S., Periyasamy-Thandavan, S., Fulzele, S., Hamrick, M.W., et al. (2019). What doesn’t kill you makes you stranger: Dipeptidyl peptidase-4 (CD26) proteolysis differentially modulates the activity of many peptide hormones and cytokines generating novel cryptic bioactive ligands. Pharmacol Ther 198, 90–108. 10.1016/j.pharmthera.2019.02.005.

48. Wilson, A.L., Moffitt, L.R., Wilson, K.L., Bilandzic, M., Wright, M.D., Gorrell, M.D., Oehler, M.K., Plebanski, M., and Stephens, A.N. (2021). DPP4 Inhibitor Sitagliptin Enhances Lymphocyte Recruitment and Prolongs Survival in a Syngeneic Ovarian Cancer Mouse Model. Cancers (Basel) 13. 10.3390/cancers13030487.

49. Fitzgerald, A.A., Wang, S., Agarwal, V., Marcisak, E.F., Zuo, A., Jablonski, S.A., Loth, M., Fertig, E.J., MacDougall, J., Zhukovsky, E., et al. (2021). DPP inhibition alters the CXCR3 axis and enhances NK and CD8+ T cell infiltration to improve anti-PD1 efficacy in murine models of pancreatic ductal adenocarcinoma. J Immunother Cancer 9. 10.1136/jitc-2021-002837.

50. Nishina, S., Yamauchi, A., Kawaguchi, T., Kaku, K., Goto, M., Sasaki, K., Hara, Y., Tomiyama, Y., Kuribayashi, F., Torimura, T., and Hino, K. (2019). Dipeptidyl Peptidase 4 Inhibitors Reduce Hepatocellular Carcinoma by Activating Lymphocyte Chemotaxis in Mice. Cell Mol Gastroenterol Hepatol 7, 115–134. 10.1016/j.jcmgh.2018.08.008.

51. Mathewson, N.D., Ashenberg, O., Tirosh, I., Gritsch, S., Perez, E.M., Marx, S., Jerby-Arnon, L., Chanoch-Myers, R., Hara, T., Richman, A.R., et al. (2021). Inhibitory CD161 receptor identified in glioma-infiltrating T cells by single-cell analysis. Cell 184, 1281–1298 e1226. 10.1016/j.cell.2021.01.022.

52. Chang, C.H., and Pearce, E.L. (2016). Emerging concepts of T cell metabolism as a target of immunotherapy. Nat Immunol 17, 364–368. 10.1038/ni.3415.

53. Bengsch, B., Seigel, B., Flecken, T., Wolanski, J., Blum, H.E., and Thimme, R. (2012). Human Th17 cells express high levels of enzymatically active dipeptidylpeptidase IV (CD26). J Immunol 188, 5438–5447. 10.4049/jimmunol.1103801.

54. Morimoto, C., Torimoto, Y., Levinson, G., Rudd, C.E., Schrieber, M., Dang, N.H., Letvin, N.L., and Schlossman, S.F. (1989). 1F7, a novel cell surface molecule, involved in helper function of CD4 cells. J Immunol 143, 3430–3439.

55. Gebhardt, T., Park, S.L., and Parish, I.A. (2023). Stem-like exhausted and memory CD8(+) T cells in cancer. Nat Rev Cancer 23, 780–798. 10.1038/s41568-023-00615-0.

56. Doering, T.A., Crawford, A., Angelosanto, J.M., Paley, M.A., Ziegler, C.G., and Wherry, E.J. (2012). Network analysis reveals centrally connected genes and pathways involved in CD8+ T cell exhaustion versus memory. Immunity 37, 1130–1144. 10.1016/j.immuni.2012.08.021.

57. Wang, H., Kadlecek, T.A., Au-Yeung, B.B., Goodfellow, H.E., Hsu, L.Y., Freedman, T.S., and Weiss, A. (2010). ZAP-70: an essential kinase in T-cell signaling. Cold Spring Harb Perspect Biol 2, a002279. 10.1101/cshperspect.a002279.

58. Badia, R., Ballana, E., Castellvi, M., Garcia-Vidal, E., Pujantell, M., Clotet, B., Prado, J.G., Puig, J., Martinez, M.A., Riveira-Munoz, E., and Este, J.A. (2018). CD32 expression is associated to T-cell activation and is not a marker of the HIV-1 reservoir. Nat Commun 9, 2739. 10.1038/s41467-018-05157-w.

59. Weil, R., and Israel, A. (2006). Deciphering the pathway from the TCR to NF-kappaB. Cell Death Differ 13, 826–833. 10.1038/sj.cdd.4401856.

60. Shah, K., Al-Haidari, A., Sun, J., and Kazi, J.U. (2021). T cell receptor (TCR) signaling in health and disease. Signal Transduct Target Ther 6, 412. 10.1038/s41392-021-00823-w.

61. Cavalcanti-de-Albuquerque, J.P., de-Souza-Ferreira, E., de Carvalho, D.P., and Galina, A. (2022). Coupling of GABA Metabolism to Mitochondrial Glucose Phosphorylation. Neurochem Res 47, 470–480. 10.1007/s11064-021-03463-2.

62. Ravasz, D., Kacso, G., Fodor, V., Horvath, K., Adam-Vizi, V., and Chinopoulos, C. (2017). Catabolism of GABA, succinic semialdehyde or gamma-hydroxybutyrate through the GABA shunt impair mitochondrial substrate-level phosphorylation. Neurochem Int 109, 41–53. 10.1016/j.neuint.2017.03.008.

63. Paine, T.A., Cooke, E.K., and Lowes, D.C. (2015). Effects of chronic inhibition of GABA synthesis on attention and impulse control. Pharmacol Biochem Behav 135, 97–104. 10.1016/j.pbb.2015.05.019.

64. Zhu, X., Li, Q., and Zhu, X. (2022). Mechanisms of CAR T cell exhaustion and current counteraction strategies. Front Cell Dev Biol 10, 1034257. 10.3389/fcell.2022.1034257.

65. Delgoffe, G.M., Xu, C., Mackall, C.L., Green, M.R., Gottschalk, S., Speiser, D.E., Zehn, D., and Beavis, P.A. (2021). The role of exhaustion in CAR T cell therapy. Cancer Cell 39, 885–888. 10.1016/j.ccell.2021.06.012.

66. Bagley, S.J., Logun, M., Fraietta, J.A., Wang, X., Desai, A.S., Bagley, L.J., Nabavizadeh, A., Jarocha, D., Martins, R., Maloney, E., et al. (2024). Intrathecal bivalent CAR T cells targeting EGFR and IL13Ralpha2 in recurrent glioblastoma: phase 1 trial interim results. Nat Med 30, 1320–1329. 10.1038/s41591-024-02893-z.

67. Bagley, S.J., Desai, A.S., Fraietta, J.A., Silverbush, D., Chafamo, D., Freeburg, N.F., Gopikrishna, G.K., Rech, A.J., Nabavizadeh, A., Bagley, L.J., et al. (2025). Intracerebroventricular bivalent CAR T cells targeting EGFR and IL-13Ralpha2 in recurrent glioblastoma: a phase 1 trial. Nat Med. 10.1038/s41591-025-03745-0.

68. Vitanza, N.A., Ronsley, R., Choe, M., Seidel, K., Huang, W., Rawlings-Rhea, S.D., Beam, M., Steinmetzer, L., Wilson, A.L., Brown, C., et al. (2025). Intracerebroventricular B7-H3-targeting CAR T cells for diffuse intrinsic pontine glioma: a phase 1 trial. Nat Med 31, 861–868. 10.1038/s41591-024-03451-3.

69. Han, J., Khatwani, N., Searles, T.G., Turk, M.J., and Angeles, C.V. (2020). Memory CD8(+) T cell responses to cancer. Semin Immunol 49, 101435. 10.1016/j.smim.2020.101435.

70. Klebanoff, C.A., Gattinoni, L., and Restifo, N.P. (2006). CD8+ T-cell memory in tumor immunology and immunotherapy. Immunol Rev 211, 214–224. 10.1111/j.0105-2896.2006.00391.x.

71. Das, S., and Johnson, D.B. (2019). Immune-related adverse events and anti-tumor efficacy of immune checkpoint inhibitors. J Immunother Cancer 7, 306. 10.1186/s40425-019-0805-8.

72. Sharma, P., Hu-Lieskovan, S., Wargo, J.A., and Ribas, A. (2017). Primary, Adaptive, and Acquired Resistance to Cancer Immunotherapy. Cell 168, 707–723. 10.1016/j.cell.2017.01.017.

73. Wu, B., Zhang, B., Li, B., Wu, H., and Jiang, M. (2024). Cold and hot tumors: from molecular mechanisms to targeted therapy. Signal Transduct Target Ther 9, 274. 10.1038/s41392-024-01979-x.

74. Brunell, A.E., Lahesmaa, R., Autio, A., and Thotakura, A.K. (2023). Exhausted T cells hijacking the cancer-immunity cycle: Assets and liabilities. Front Immunol 14, 1151632. 10.3389/fimmu.2023.1151632.

75. Shyer, J.A., Flavell, R.A., and Bailis, W. (2020). Metabolic signaling in T cells. Cell Res 30, 649–659. 10.1038/s41422-020-0379-5.

76. de Aquino, M.T.P., Hodo, T.W., Ochoa, S.G., Uzhachenko, R.V., Mohammed, M.A., Goodwin, J.S., Kanagasabai, T., Ivanova, A.V., and Shanker, A. (2025). Glutamate receptor-T cell receptor signaling potentiates full CD8(+) T cell activation and effector function in tumor immunity. iScience 28, 112772. 10.1016/j.isci.2025.112772.

77. Carr, E.L., Kelman, A., Wu, G.S., Gopaul, R., Senkevitch, E., Aghvanyan, A., Turay, A.M., and Frauwirth, K.A. (2010). Glutamine uptake and metabolism are coordinately regulated by ERK/MAPK during T lymphocyte activation. J Immunol 185, 1037–1044. 10.4049/jimmunol.0903586.

78. Morris, G., Gevezova, M., Sarafian, V., and Maes, M. (2022). Redox regulation of the immune response. Cell Mol Immunol 19, 1079–1101. 10.1038/s41423-022-00902-0.

79. Bailey, S.R., Nelson, M.H., Majchrzak, K., Bowers, J.S., Wyatt, M.M., Smith, A.S., Neal, L.R., Shirai, K., Carpenito, C., June, C.H., et al. (2017). Human CD26(high) T cells elicit tumor immunity against multiple malignancies via enhanced migration and persistence. Nat Commun 8, 1961. 10.1038/s41467-017-01867-9.

80. Nelson, M.H., Knochelmann, H.M., Bailey, S.R., Huff, L.W., Bowers, J.S., Majchrzak-Kuligowska, K., Wyatt, M.M., Rubinstein, M.P., Mehrotra, S., Nishimura, M.I., et al. (2020). Identification of human CD4(+) T cell populations with distinct antitumor activity. Sci Adv 6. 10.1126/sciadv.aba7443.

81. Cordero, O.J., Yang, C.P., and Bell, E.B. (2007). On the role of CD26 in CD4 memory T cells. Immunobiology 212, 85–94. 10.1016/j.imbio.2006.12.002.

82. Yang, Q., Fu, B., Luo, D., Wang, H., Cao, H., Chen, X., Tian, L., and Yu, X. (2022). The Multiple Biological Functions of Dipeptidyl Peptidase-4 in Bone Metabolism. Front Endocrinol (Lausanne) 13, 856954. 10.3389/fendo.2022.856954.

83. Ng, II, Zhang, J., Tian, T., Peng, Q., Huang, Z., Xiao, K., Yao, X., Ng, L., Zeng, J., and Tang, H. (2024). Network-based screening identifies sitagliptin as an antitumor drug targeting dendritic cells. J Immunother Cancer 12. 10.1136/jitc-2023-008254.

84. Hollande, C., Boussier, J., Ziai, J., Nozawa, T., Bondet, V., Phung, W., Lu, B., Duffy, D., Paradis, V., Mallet, V., et al. (2019). Inhibition of the dipeptidyl peptidase DPP4 (CD26) reveals IL-33-dependent eosinophil-mediated control of tumor growth. Nat Immunol 20, 257–264. 10.1038/s41590-019-0321-5.

85. Pinheiro, M.M., Stoppa, C.L., Valduga, C.J., Okuyama, C.E., Gorjao, R., Pereira, R.M., and Diniz, S.N. (2017). Sitagliptin inhibit human lymphocytes proliferation and Th1/Th17 differentiation in vitro. Eur J Pharm Sci 100, 17–24. 10.1016/j.ejps.2016.12.040.

86. Aso, Y., Fukushima, M., Sagara, M., Jojima, T., Iijima, T., Suzuki, K., Momobayashi, A., Kasai, K., and Inukai, T. (2015). Sitagliptin, a DPP-4 inhibitor, alters the subsets of circulating CD4+ T cells in patients with type 2 diabetes. Diabetes Res Clin Pract 110, 250–256. 10.1016/j.diabres.2015.10.012.

87. Chao, R., Nishida, M., Yamashita, N., Tokumasu, M., Zhao, W., Kudo, I., and Udono, H. (2022). Nutrient Condition in the Microenvironment Determines Essential Metabolisms of CD8(+) T Cells for Enhanced IFNgamma Production by Metformin. Front Immunol 13, 864225. 10.3389/fimmu.2022.864225.

88. Ton Nu, Q.C., and Park, P.H. (2025). Metabolic modulation of CD8(+) T cells by metformin: A promising adjuvant strategy for CD8(+) T Cell-Based Immunotherapies. Pharmacol Res 222, 108015. 10.1016/j.phrs.2025.108015.

89. Finisguerra, V., Dvorakova, T., Formenti, M., Van Meerbeeck, P., Mignion, L., Gallez, B., and Van den Eynde, B.J. (2023). Metformin improves cancer immunotherapy by directly rescuing tumor-infiltrating CD8 T lymphocytes from hypoxia-induced immunosuppression. J Immunother Cancer 11. 10.1136/jitc-2022-005719.

90. Ali, A., Fuentes, A., Skelton, W.I., Wang, Y., McGorray, S., Shah, C., Bishnoi, R., Dang, L.H., and Dang, N.H. (2019). A multi-center retrospective analysis of the effect of DPP4 inhibitors on progression-free survival in advanced airway and colorectal cancers. Mol Clin Oncol 10, 118–124. 10.3892/mco.2018.1766.

91. Ng, L., Foo, D.C., Wong, C.K., Man, A.T., Lo, O.S., and Law, W.L. (2021). Repurposing DPP-4 Inhibitors for Colorectal Cancer: A Retrospective and Single Center Study. Cancers (Basel) 13. 10.3390/cancers13143588.

92. Lee, J.H., Kim, T.I., Jeon, S.M., Hong, S.P., Cheon, J.H., and Kim, W.H. (2012). The effects of metformin on the survival of colorectal cancer patients with diabetes mellitus. Int J Cancer 131, 752–759. 10.1002/ijc.26421.

93. Haydar, D., Houke, H., Chiang, J., Yi, Z., Ode, Z., Caldwell, K., Zhu, X., Mercer, K.S., Stripay, J.L., Shaw, T.I., et al. (2021). Cell-surface antigen profiling of pediatric brain tumors: B7-H3 is consistently expressed and can be targeted via local or systemic CAR T-cell delivery. Neuro Oncol 23, 999–1011. 10.1093/neuonc/noaa278.

94. Jacobson, G., Cicchiello, A., Sutton, J.P., and Shah, A. (2021). Medicare Advantage vs. Traditional Medicare: How Do Beneficiaries’ Characteristics and Experiences Differ? Commonwealth Fund. October 2021. 10.26099/yxq0-1w42.

95. Motheral, B.R., and Fairman, K.A. (1997). The use of claims databases for outcomes research: rationale, challenges, and strategies. Clin. Ther. 19, 346–366. 10.1016/S0149-2918(97)80122-1.

96. Strom, B.L., and Carson, J.L. (1990). Use of automated databases for pharmacoepidemiology research. Epidemiol. Rev. 12, 87–107. 10.1093/oxfordjournals.epirev.a036064.

97. Jollis, J.G., Ancukiewicz, M., DeLong, E.R., Pryor, D.B., Muhlbaier, L.H., and Mark, D.B. (1993). Discordance of databases designed for claims payment versus clinical information systems. Implications for outcomes research. Ann. Intern. Med. 119, 844–850. 10.7326/0003-4819-119-8-199310150-00011.

98. Strom, B.L. (2001). Data validity issues in using claims data. Pharmacoepidemiol Drug Saf 10, 389–392. 10.1002/pds.610.

99. SEER-Medicare Linked Database. (2022).

100. Price, M., Ballard, C., Benedetti, J., Neff, C., Cioffi, G., Waite, K.A., Kruchko, C., Barnholtz-Sloan, J.S., and Ostrom, Q.T. (2024). CBTRUS Statistical Report: Primary Brain and Other Central Nervous System Tumors Diagnosed in the United States in 2017-2021. Neuro Oncol. 26, vi1–vi85. 10.1093/neuonc/noae145.

101. Healthcare Delivery Research Program (2021). Comorbidity SAS Macro (2021 version). https://healthcaredelivery.cancer.gov/seermedicare/considerations/macro-2021.html.

102. Yu, K., Hu, Y., Wu, F., Guo, Q., Qian, Z., Hu, W., Chen, J., Wang, K., Fan, X., Wu, X., et al. (2020). Surveying brain tumor heterogeneity by single-cell RNA-sequencing of multi-sector biopsies. Natl Sci Rev 7, 1306–1318. 10.1093/nsr/nwaa099.

103. Caruso, F.P., Garofano, L., D’Angelo, F., Yu, K., Tang, F., Yuan, J., Zhang, J., Cerulo, L., Pagnotta, S.M., Bedognetti, D., et al. (2020). A map of tumor-host interactions in glioma at single-cell resolution. Gigascience 9. 10.1093/gigascience/giaa109.

104. van Dijk, D., Sharma, R., Nainys, J., Yim, K., Kathail, P., Carr, A.J., Burdziak, C., Moon, K.R., Chaffer, C.L., Pattabiraman, D., et al. (2018). Recovering Gene Interactions from Single-Cell Data Using Data Diffusion. Cell 174, 716–729 e727. 10.1016/j.cell.2018.05.061.

105. Frattini, V., Pagnotta, S.M., Tala, Fan, J.J., Russo, M.V., Lee, S.B., Garofano, L., Zhang, J., Shi, P., Lewis, G., et al. (2018). A metabolic function of FGFR3-TACC3 gene fusions in cancer. Nature 553, 222–227. 10.1038/nature25171.

